# The impact of patient biology on racial disparities in breast cancer outcome

**DOI:** 10.1101/2025.11.05.685148

**Authors:** JT DeWitt, D Jimenez-Tovar, C Nguyen, E Oropeza, M Raghunathan, ET Karabay, A Lamichhane, E Kirk, SA Raghavan, P Katira, C Luna Lopez, MA Troester, SJ Freedland, S Haricharan

## Abstract

Hormone receptor positive (HR+) breast cancer is the most common subtype of breast cancer diagnosed globally. Despite effective targeted therapies, HR+ breast cancer remains a leading cause of cancer-related death in women. Long-standing epidemiological research identifies significantly worse outcomes for Black women diagnosed with HR+ breast cancer relative to White women. While structural factors such as access to healthcare and education level contribute to this outcome disparity, it persists even in analyses where these factors are controlled. In-depth analyses of the somatic molecular biology that may underlie these outcome disparities are hampered by a lack of datasets that represent Black patient populations. Here, we generate a HR+ breast cancer patient transcriptomic dataset that overrepresents Black women and controls for access to healthcare and education level. We find that signatures relating to the tumor microenvironment, i.e. collagen deposition and prognostically unfavorable T-cell landscapes are enriched in HR+ tumors from Black women. Importantly, we find, using experimental model systems *in vitro* and *in vivo*, that race-aligned collagen deposition patterns are at least partially attributable to tumor cell-intrinsic signaling and critical for Black breast cancer metastasis. We also find that unfavorable T-cell signatures in HR+ tumors from Black women, which have previously been attributed to race and ancestry, are more strongly poverty-aligned. Using multiple independent datasets, we identify STAT4 as a potential master regulator of this poverty-associated tumor immune signature. Together, these findings provide new evidence that somatic molecular biology of breast cancer patients can be modified by multiple structural factors such as self-identified race and poverty burden to promote poor patient outcomes. Integrating an understanding of structural factors into molecular cancer research is critical for implementing truly personalized, and maximally effective, oncology systems.

## Introduction

Despite striking advances in technology and better access to patient data, precision diagnostics and therapeutics remain unobtainable for many cancer patients. Cancer datasets and experimental model systems overrepresent the disease in affluent, non-Hispanic White (henceforth, referred to as White) patients, while typically underrepresenting all other patient demographics^1,2^. Perhaps unsurprisingly, there is a widening and persistent gap in cancer survivorship between affluent, White patients and other racial and ethnic groups (reviewed in^3,4^). Specifically, non-Hispanic Black (henceforth, termed Black) and Hispanic cancer patients have worse outcomes than White counterparts across many different cancer types, including hormone receptor positive (HR+) breast cancer^4–6^.

HR+ breast cancer is the most common breast cancer diagnosis, affecting over 240,000 people each year, in the US alone^4^, and is a leading cause of cancer-related death in women. Black women diagnosed with HR+ breast cancer have a 40% higher likelihood of dying from their diagnosis than White women^4,7^. Some of the factors contributing to this worsened outcome are structural including poor access to healthcare and lower education levels, which disproportionately affect Black people in the US^8,9^. However, even when these factors are controlled for, the disparity in HR+ breast cancer outcome largely persists^7,10^, suggesting that a multitude of patient-intrinsic and -extrinsic factors are likely involved^11,12^.

Studies of a role for genetic ancestry in breast cancer outcome disparities typically focus on the triple negative subtype, which women of African descent are nearly twice as likely to be diagnosed with compared to women of European descent^13^. This is indeed a valid and important area of research. However, despite increased incidence of triple negative breast cancer in women of African descent, HR+ breast cancer remains the most common diagnosis, and cause of breast cancer-related death in both Black and White women^4^. Because HR+ breast cancer in Black women is understudied, there is a significant gap in understanding whether the somatic molecular biology of HR+ tumors from Black women is distinct from that of White women. Several studies suggest that much of the 40% increase in cancer-related deaths that Black women face overall is driven by HR+, rather than triple negative, breast cancer diagnoses^7,8,10,14,15^.

Analyses of transcriptomics-based patient datasets uniformly identify unique signatures in the somatic, molecular biology of breast cancer in Black women^6,13,16–18^. However, most of these previous analyses use datasets which lack annotation of poverty burden, and other social or structural determinants of health, precluding attempts at deconvoluting the impact of race vs structural factors on outcome-relevant somatic tumor biology^19^. Epidemiological studies investigating outcome disparities, which often do incorporate such annotations, typically combine different breast cancer subtypes in their analyses to compensate for sample size and/or do not contain additional molecular biological data from patient tumors (i.e. transcriptomic, genomic, proteomic data)^19^.

Despite these intersectional caveats, there is evidence to suggest that the somatic molecular biology of Black breast cancer is distinct: characterized by an exhausted T-cell immune signature^16,20^, a distinct DNA damage repair signature^17,18^, and high proliferation/cell cycle dysregulated signatures^17,18,21^, all of which associate with worse prognosis. The extent to which these somatic molecular differences in tumor biology are impacted by extrinsic factors (such as poverty burden) in addition to ancestry, and whether they further influence phenotypes of proliferation and metastasis that underlie poor patient outcomes remains to be investigated.

Here, we address some of these questions by generating a transcriptomic dataset of 56 HR+ breast tumors that is controlled for access to healthcare, overrepresents Black women, and includes annotation of factors reflecting life experiences: education level, body-mass index (BMI), and neighborhood-level poverty indices. Analyses of these data reveal new insights into the impact of sociodemographic factors on the somatic tumor microenvironment, validated in independent datasets and in experimental model systems, as detailed below.

## Results

### Hormone receptor positive breast tumors from Black women have distinct somatic, molecular signatures

We obtained 64 HR+ primary tumors from a Veterans Affairs (VA) archival tissue bank; the majority of these tumors had corresponding clinicopathological annotations for tumor subtype, receptor status, patient outcome, and patient demographics data: poverty burden indicator, highest education level, BMI, and self-reported race (n=60) (Table 1 and 2). Fifty-six HR+ primary tumors passed quality checks and contained sufficient sample for bulk RNA sequencing and were controlled for clinicopathological characteristics and patient demographics between tumors from Black and White patients (Table 1 and 2). RNA extracted from paraffin-embedded tumor samples from these 56 patients was subsequently sequenced to create a bulk transcriptomics dataset (see REMARK diagram in Fig 1A).

**Figure 1.**
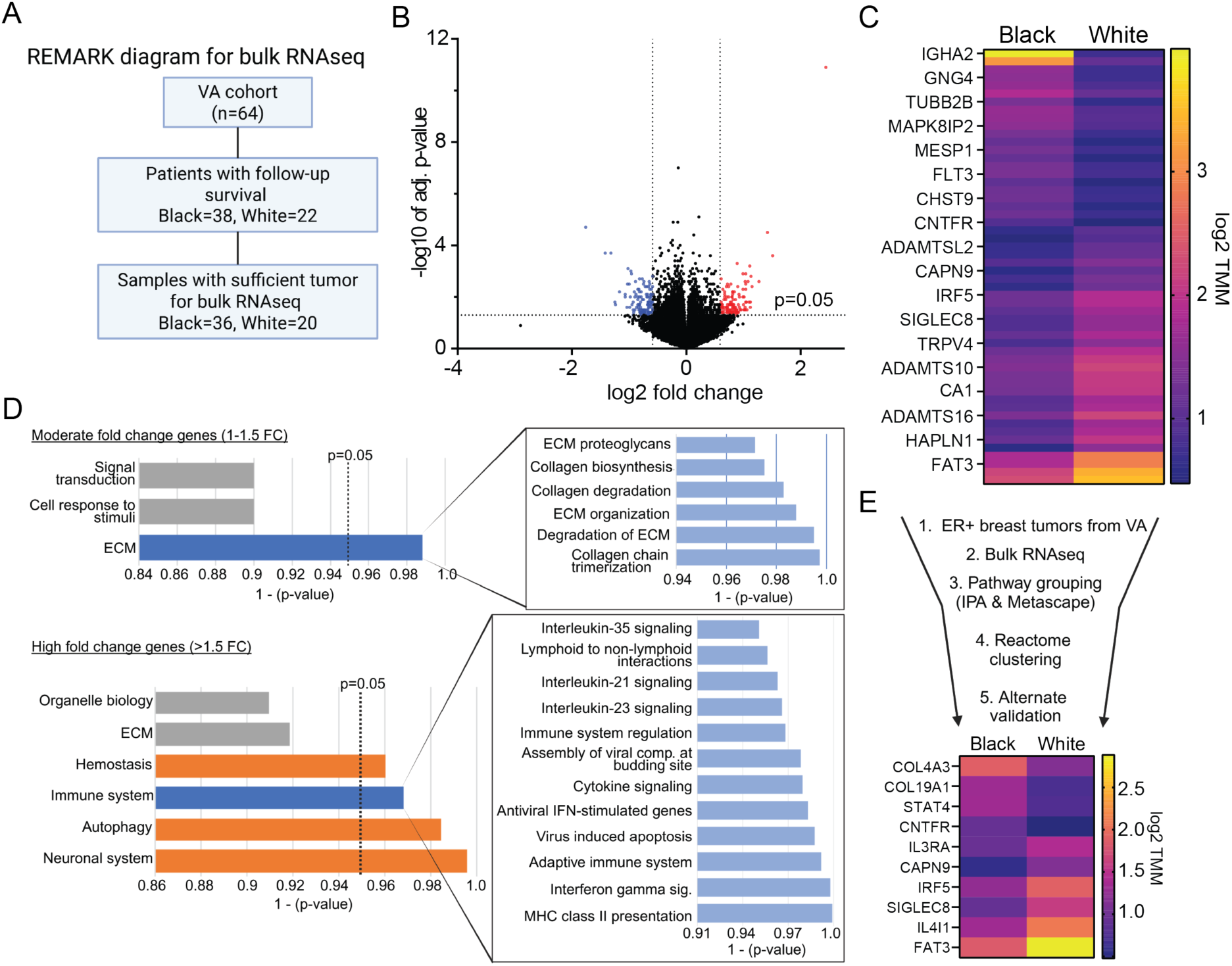
Hormone receptor positive breast tumors from Black women have distinct somatic, transcriptomic signatures than those from White women. (A) REMARK diagram of selection of archival patient tumors for bulk RNAseq analysis. (B) Volcano plot representing global transcriptomic differences in the tumors from Black women compared to White women. Blue dots indicate significantly downregulated gene expression, while red represents significant upregulation. (C) Heat-map for all significantly differently (p<0.05) expressed RNA candidates in tumor from Black women compared to White women. (D) Summary plots from Reactome pathway grouping genes with moderate fold change (1-1.5), and high fold change (>1.5) difference between tumors from Black and White women. Associated GSEA data in Fig S1. (E) Funnel schematic detailing the pipeline for identifying and validating differently regulated candidates, and a heat-map representing differences in gene expression between Black and White tumors for all candidates that validate in at least one other independent dataset (TCGA and Ellsworth). Abbreviations: VA (Veterans Affairs), adj. (adjusted), TMM (Trimmed Mean of M values), ECM (extracellular matrix), FC (fold change), and IPA (Ingenuity Pathway Analysis).

**Table 1.**
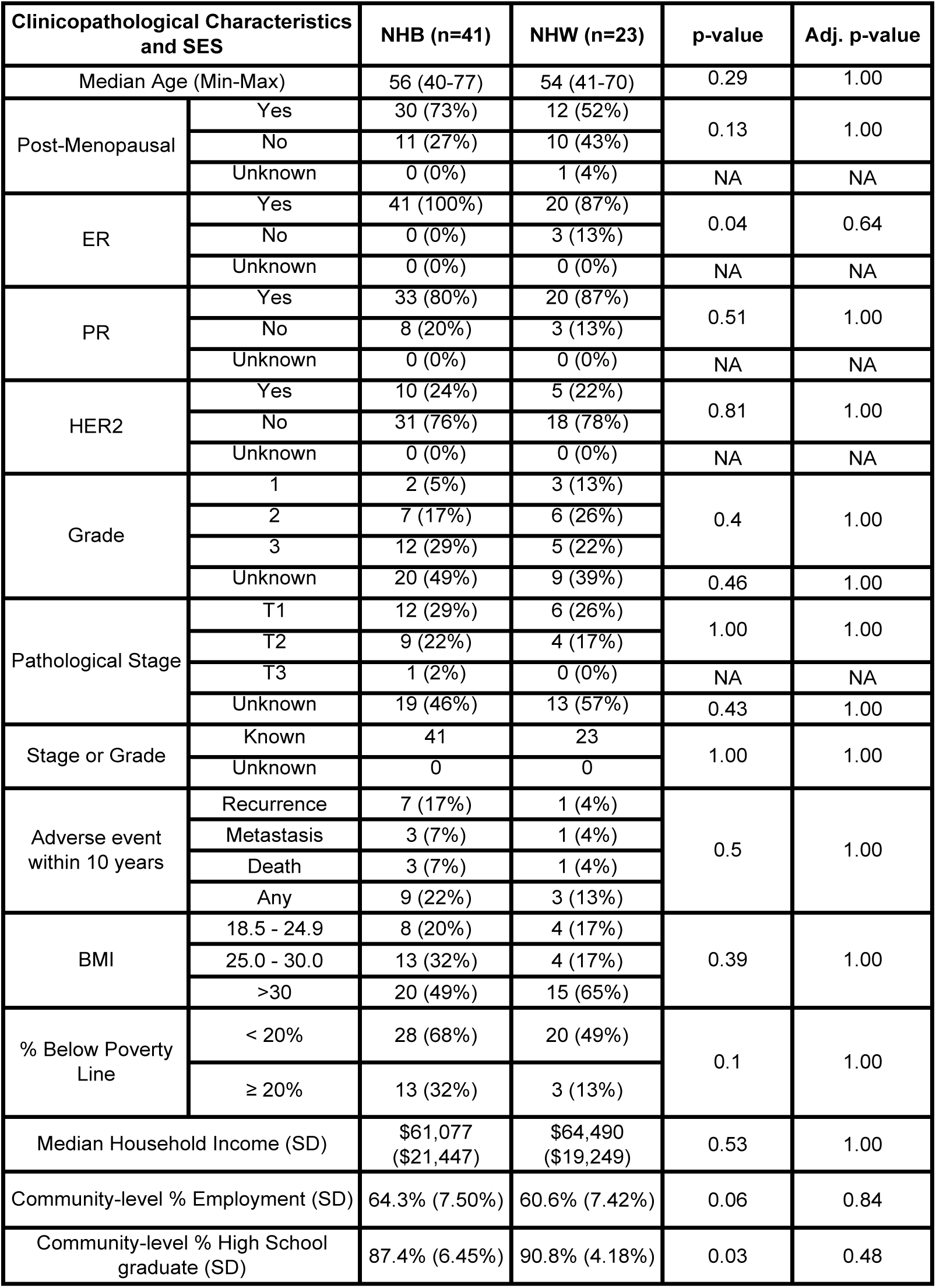
Clinicopathological characteristics for the patient cohort from Veterans Affairs. In this dataset, all patients were diagnosed and treated within the Veterans Affairs healthcare systems in Durham, North Carolina. BI-RADS scoring was performed prior to tumor collection by core needle biopsies or fine needle aspiration to receive a pathological tumor grade. All patients received either pathological stage, or grade. An ‘Unknown’ designation indicates data is not reported in annotations. Students’ T-tests were used to determine p-values for continuous data and by chi-squared for categorical tests. Adjusted p-values were determined by Holms-Bonferroni correction. Pathological stage was identified by TNM classification. NA= not available, adj. p-value= adjusted p-value.

**Table 2.** Systemic metrics of structural factors reflecting life experiences for the patient cohort from Veterans Affairs. Body-mass index (BMI) metrics were reported for patients in the VA dataset. Poverty burden indices, percent employment, and percent high school graduates are reported as community-level percentages using geolocation. For poverty burden, percentages reflect the proportion of people living below the Federal poverty line in that zip code. Students’ T-tests were used to determine p-values. Adjusted p-values were determined by Holms-Bonferroni correction. NA= not available, adj. p-value= adjusted p-value.

| Socioeconomic factors |  | Black (n=41) | White (n=23) | p-value | Adj. p-value |
| --- | --- | --- | --- | --- | --- |
|  |  | n (%) |  |  |  |
| BMI | 18.5 - 24.9 | 8 (20%) | 4 (17%) | 0.39 | 0.49 |
|  | 25.0 - 30.0 | 13 (32%) | 4 (17%) |  |  |
|  | >30 | 20 (49%) | 15 (65%) |  |  |
| % Below Poverty Line | < 20% | 28 (68%) | 20 (49%) | 0.1 | 0.17 |
|  | ≥ 20% | 13 (32%) | 3 (13%) |  |  |
| Socioeconomic factors |  | Black (n=41) | White (n=23) | p-value | Adj. p-value |
|  |  | median (SD) |  |  |  |
| Household Income | | \$61,077<br>(\$21,447) | \$64,490<br>(\$19,249) | 0.53 | 0.53 |
| Community-level % Employment |  | 64.3% (7.50%) | 60.6% (7.42%) | 0.06 | 0.15 |
| Community-level % High School graduate |  | 87.4% (6.45%) | 90.8% (4.18%) | 0.03 | 0.15 |

Comparison of gene expression between the transcriptome of Black (n=36) and White (n=20) patient tumor samples from this dataset identified many differences between the two groups despite controlled clinicopathological characteristics and access to healthcare (Fig 1B-C). Specifically, we found significant enrichment in extracellular matrix (ECM) and immune signatures in tumors from Black women (Fig 1D; Metascape and IPA analyses presented in Fig S1A). Genes with high fold change differences (>1.5 FC) map to immune pathways, and genes with moderate-fold differences (1-1.5 FC) to ECM pathways (Fig 1D & Reacfoam in Fig S1B). We then cross-validated genes that are significantly differently expressed in tumors from Black vs White women in this VA dataset, for similar differences between these two patient demographics in two independent public RNA sequencing datasets: TCGA^22^ and Ellsworth^23^. Of the 54 candidate genes identified as significantly differently expressed in this VA dataset, 10 validated as being differently expressed in HR+ tumors from Black women in at least one independent dataset (Fig 1E). Validated candidate genes predominately relate to ECM and immune pathways emphasizing the importance of these two pathways in describing transcriptomic features unique to HR+ tumors in Black women.

### Black women have tumor-intrinsic signaling that induces dense, radial organization of collagen in the tumor microenvironment

Twenty percent (2/10) of candidate genes validated as differently expressed in tumors from Black patients relative to those from White patients are directly linked to collagen deposition and organization (*COL4A3* and *COL19A1*) and an additional forty percent (4/10) candidate genes are linked to collagen regulation (*CNTFR*, *IL3RA*, *IRF5* and *FAT3*). Therefore, we conducted Masson’s Trichrome on a subset of patient tumors from the VA dataset with sufficient sample remaining following bulk RNAseq (Total: n=23, Black: n=14, White: n=9) to quantify collagen abundance and organization (Fig 2A). Overall, we detected higher density of deposited collagen in tumors from Black women than those from White women (2.3-fold increase in tumoral collagen; p= 0.0364) (Fig 2B). Orthogonal, automated quantification of collagen by second harmonic generation confirmed this observation (2.7-fold increase in tumoral collagen density; p= 3.7 x10^-9^) (Fig 2C). Collagen fibers surrounding tumoral ducts are also differently organized in tumors from Black women (Fig 2D). Specifically, collagen fibers in tumors from Black women increasingly align at radial angles (90⁰-120⁰ relative to the horizontal plane) while those in tumors from White women are more evenly distributed across all angles with no apparent enrichment across any angular distribution (p=0.014) (Fig 2E). Prior investigations have reported inverse correlation between BMI and breast tissue density in breast cancer patients^24^, however, we did not observe such correlation in this VA cohort (All: R^2^=0.04, p=0.35; Black: R^2^=0.0002, p=0.96; White: R^2^=0.17, p=0.23) (Fig S2A).

**Figure 2.**
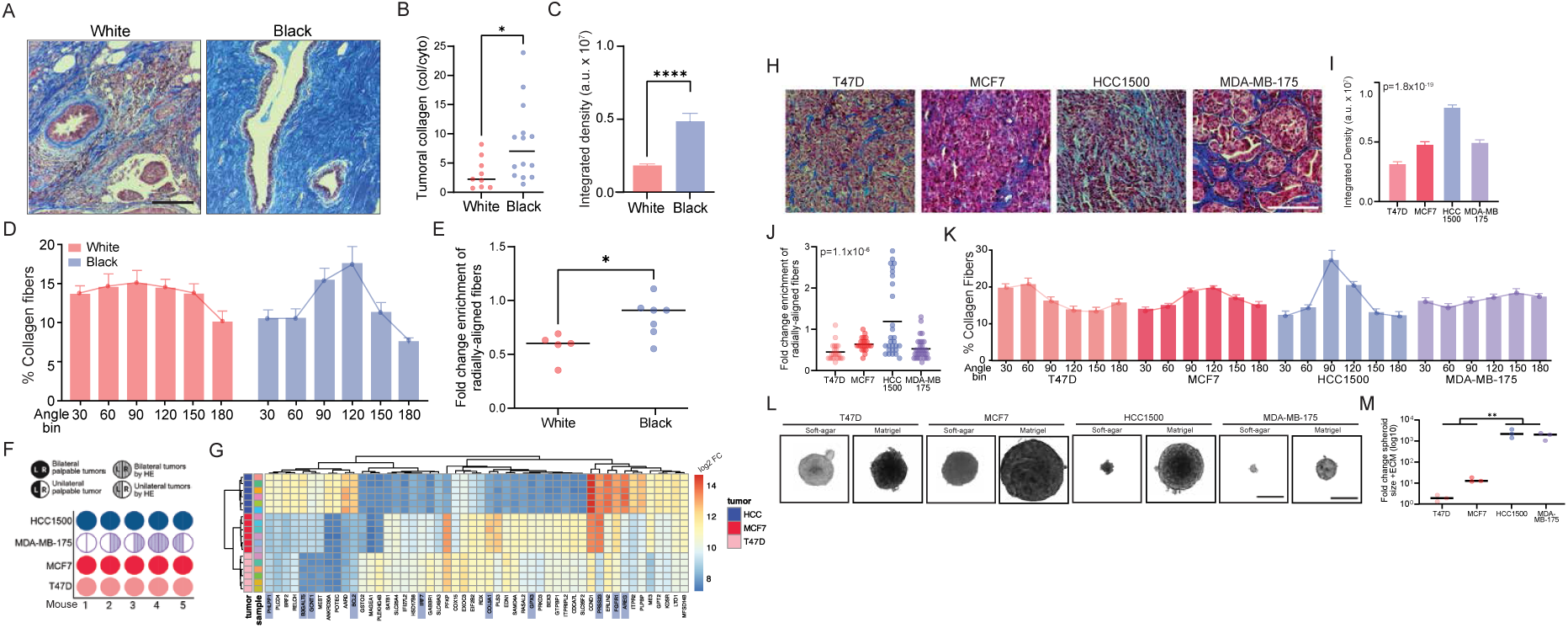
Hormone receptor positive breast tumors from Black women have differently deposited and organized collagen compared to those from White women. (A, H) Representative trichrome stains for breast tumor sections from Black and White women (A) and from tumors from cell lines derived from Black (HCC-1500, MDA-MB-175) and White (MCF7, T47D) women xenografted into fat pads of nude mice (H). (B) Dot-plot for FiJi quantifications of collagen levels relative to cellular content, horizontal line represents average collagen content. (C-E, I-K) Collagen quantifications by second harmonic generation showing integrated collagen density (C, I), average percent collagen fibers by angle binning (D, K), and fold change enrichment for radially-aligned collagen fibers between 90-120° relative to tumor cell reference with black line at average (E, J). (F) Tumor cell xenograft take-rates from cell lines derived from Black (HCC1500 and MDA-MB-175) and White (MCF7 and T47D) breast cancer patients and injected into the fat pads of nude mice. (G) RNA sequencing heat-map of top 50 differentially regulated genes in xenograft tumors. Supporting RNAseq analysis in Fig S2. Blue shading indicates genes with known direct and indirect roles involved in collagen or the extracellular matrix. (L-M) 3D spheroids of tumor cells from HR+ breast tumor cells from Black or White women grown with (matrigel) and without (soft-agar) addition of extracellular matrix with accompanying quantifications represented as dot plots (M). Scale bars are 200um (A), 170um (H) and 350um (L). Student’s t-test (B-D, L), and one-way ANOVA (H) were used to determine p-values. Error bars depict standard deviation for all bar graphs. Statistically significant p values were graphically presented as asterisks, with one asterisk (p<0.05), two asterisks (p<0.01), three asterisks (p<0.001), and four asterisks (p<0.0001). Abbreviations: a.u. (arbitrary units), ns (not significant), L (left fat-pad), R (right fat-pad), HE (Hematoxylin and Eosin).

To test whether HR+ breast cancer cells from Black women regulate collagen deposition in their microenvironment through tumor cell-intrinsic mechanisms, we generated tumor xenografts from a panel of HR+ breast cancer cell lines. Two lines derived from Black patients (HCC1500 and MDA-MB-175) and two from White (MCF7 and T47D) were grown in mammary fat pads of nude mice. One of the lines derived from Black women, MDA-MB-175, had a significantly lower take rate than the other lines (p=0.0471) (Fig 2F) as assessed by emergence of palpable tumors. However, histological assessment of mammary fat pads injected with MDA-MB-175 cells (Fig S2B) identified sub-palpable soft-textured tumors with 60% take rate (Fig 2F). For all tumor xenografts, other than MDA-MB-175, we were able to consistently measure tumor volume. We found differences in baseline growth, but all three lines achieved exponential growth and generated defined tumors at the time of harvest (Fig S2C-D). Bulk transcriptomic analysis using RNAseq of the resultant tumors found that gene expression signatures can distinguish tumors from each cell line in a principal component analysis (Fig S2E). Moreover, these tumors reflect enrichment in gene expression signatures related to cell cycle and apoptosis in HCC1500 relative to cell lines originating from White women, which is in accordance with previously published literature from patient tumors^17–19^ (Fig S2F-G). Importantly, these tumors also reflect differences in collagen and ECM-related genes between HCC1500 and the lines originating from White women (Fig 2G). In fact, four of the top differently enriched gene expression signatures from Reactome analysis of HCC1500 tumors compared to MCF7, although not T47D, tumors relate to collagen organization and ECM (Fig S2F-G).

We next used Masson’s Trichrome as before to test whether xenograft tumors resulting from cell lines derived from Black breast cancer patients can alter collagen organization through tumor-intrinsic mechanisms (Fig 2H). Overall, collagen density is significantly higher in HCC1500 xenograft tumors than in tumors from either of the lines originating from White women (HCC1500 vs. MCF7, FC= 1.663, p= 6.15 x 10^-11^; HCC1500 vs. T47D, FC= 2.536, p= 4.39 x 10^-18^), replicating the high collagen density phenotype detectable in tumors from Black patients (Fig 2I). Moreover, the collagen fiber alignment in the HCC1500 tumors is ∼2-fold enriched for radial alignment compared to tumors from MCF7 and T47D, also recapitulating what we observe in Black patient tumors (HCC1500 vs. MCF7, p=0.0014; HCC1500 vs. T47D, p= 8 x 10^-6^) (Fig 2J-K). On the other hand, MDA-MB-175 tumors are unable to replicate either the increased tumoral collagen density (MDA-MB-175 vs. MCF7, FC= 1.032, p= 0.7041; MDA-MB-175 vs. T47D, FC= 1.574, p= 1 x 10^-5^) (Fig 2I), or the radial enrichment of collagen organization (Fig 2J-K) solely through tumor cell-intrinsic signaling. It is striking that MDA-MB-175 cells are also unable to generate palpable tumors in xenograft experiments.

To test whether collagen is required for viable cell growth and colony formation in MDA-MB-175, we examined spheroid formation across two commonly used matrices for 3D spheroid growth: a soft agar matrix without collagen, and a collagenous matrigel. Spheroids derived from both cell lines from Black women form significantly smaller colonies in the soft agar (no collagen) matrix compared to cell lines from White women (FC= 0.0022, p= 0.0043) (Fig 2L, S3A). While the collagenous matrigel-containing matrices significantly promote colony growth and formation in all examined HR+ breast cancer cell lines (Fig 2L, S3B-D), the effect-size of growth promotion provided by a collagenous matrix is 259 times higher for Black, than White, cell lines (p=0.0021) (Fig 2M).

Together, these data suggest that collagen deposition and organization in the HR+ tumor microenvironment is at least partly driven by cell-intrinsic molecular events, can be replicated in experimental model systems *in vivo*, and may be critical for the growth of HR+ breast cancer in Black women.

### Collagen density and organization in Black women is critical for metastatic phenotypes

Collagen deposition and organization are established modulators of cancer cell migration and invasion^25,26^. Therefore, we next tested whether collagen patterns observed in HR+ tumors from Black women may contribute to tumor cell invasion using *in silico* simulation for predicting tumor cell invasion potential based on bulk RNAseq data from patient tumors (Supplemental Videos). Our simulations predict that tumor cells from Black women have low invasion potential in low collagen density that models the tumor microenvironment in White patients (Fig 3A). However, increasing collagen density to levels comparable to that seen in the microenvironment of Black patients significantly increases their invasion potential (Black 1mg/ml vs Black 2mg/ml FC= 1.926, p= 0.0009; Black 1mg/ml vs. Black 4mg/ml FC= 1.718, p =0.0203). Conversely, HR+ breast cancer cells from White women are not predicted to become more invasive with increasing collagen density but instead are predicted to become significantly less invasive (White 1mg/ml vs. White 2mg/ml FC= 00.4849, p= 0.0101; White 1mg/ml vs. White 4mg/ml FC= 0.0289, p= 5.26 x 10^-9^). We also simulated collagen fiber orientation to replicate alignment patterns seen in White and Black patient tumors (Fig 3B). In lower density simulations (1mg/ml collagen) increasing fiber alignment is predicted to increase tumor cell invasion for the White relative to the Black patient group (White vs Black 1mg/ml, Al: 0.4, p= 0.0159)). However, in high collagen density (2-4mg/ml) simulations increasing fiber alignment predicts a significant boost to tumor cell invasion for the Black but not White patient groups (Black 4mg/ml, Al: 0.0 vs Al: 0.4, p= 0.0002; White p= 0.9377).

**Figure 3.**
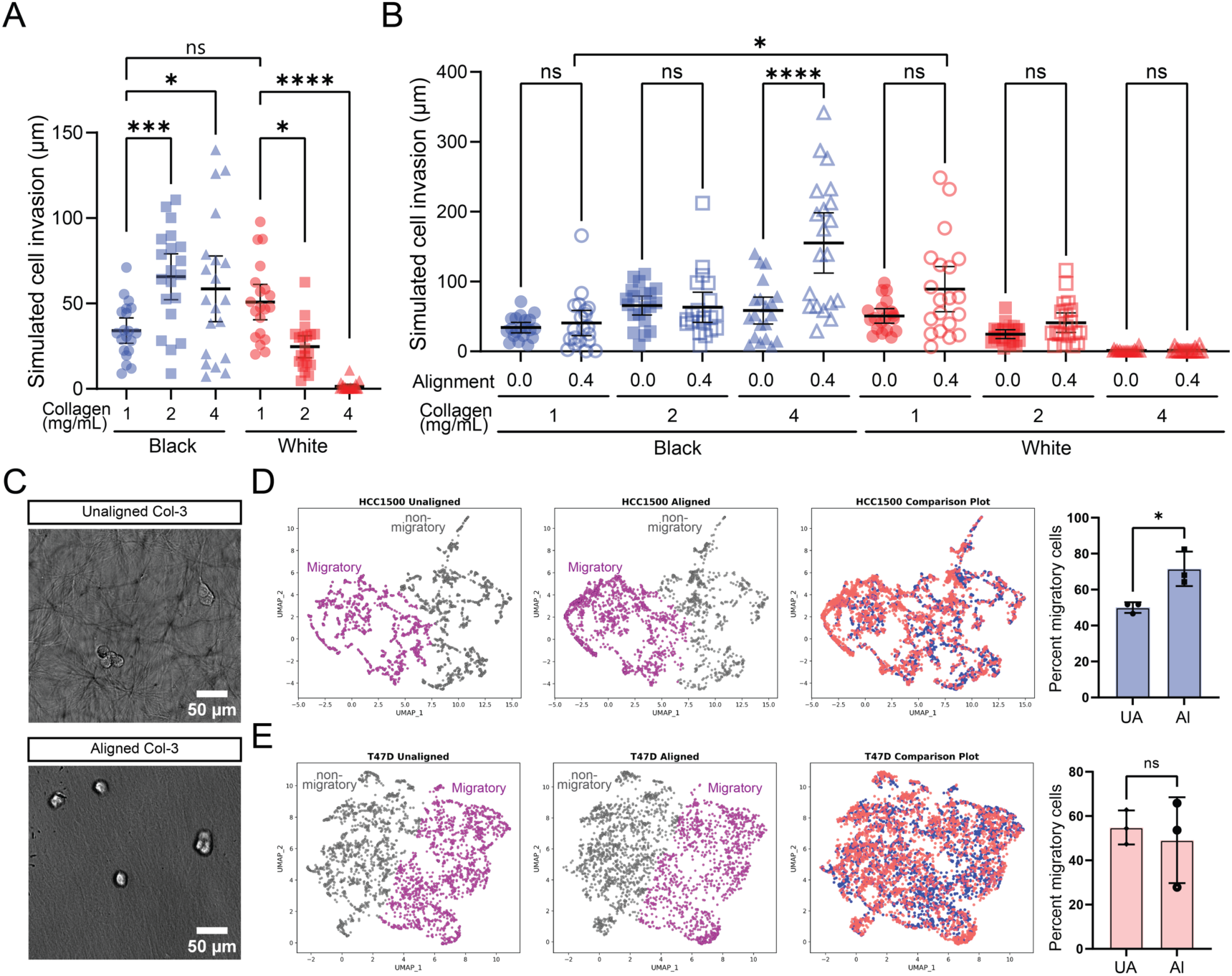
Tumoral collagen has a race-aligned impact on cancer cell migration. (A-B) Dot plots representing simulated cancer cell invasion based on bulk RNAseq data from tumors from Black and White women across (A) increasing levels of collagen deposition and (B) increasing collagen fiber alignment. (C) Representative images of unaligned and aligned collagen 1 at 3mg/mL concentrations using brightfield microscopy. (D-E) UMAP clusters for HCC1500 (D) and T47D (E) cells cultured in either aligned or unaligned collagen conditions where purple highlights migratory populations of cells and grey indicated non-migratory cell populations with the accompanying box-plot quantifications. Scale bars are 50um (C). Student’s t-test was used to determine p-values (E-F), and one-way ANOVA was used with Tukey’s test for multiple comparisons to determine adjusted p-values between groups (A-B). Statistically significant p values are graphically presented as asterisks, with one asterisk (p<0.05), two asterisks (p<0.01), three asterisks (p<0.001), and four asterisks (p<0.0001). Abbreviations: um (micron), ns (not significant), mg/mL (milligram/milliliter), ECM (extracellular matrix), Col-3 (collagen 1 at 3 mg/mL), UA (unaligned collagen), Al (aligned collagen).

Therefore, we next evaluated whether the predictions of these simulations validate experimentally using cell lines from Black or White women. To do this, we plated HCC1500 and T47D cells in two distinct high-density collagen matrices (3 mg/ml) composed of either unaligned or aligned collagen fibers (Fig 3C). We then conducted digital holographic microscopy to compare cell migratory phenotypes (Fig S3E), as measured by four morphological features: area, optical volume, eccentricity and irregularity^27^, across both collagen matrices (Fig S3F). From these parameters, we identify four unique cellular clusters in each cell line, two of which measure high for eccentricity and irregularity, suggesting increased migratory potential (Fig S3F). Mirroring the *in silico* predictions, we find that an aligned high-density collagen matrix increases the proportion of cells conforming to clusters associated with high migrator potential in the HCC1500 (p=0.049) (Fig 3D), but not in the T47D (p=0.67) (Fig 3E), lines. These findings are consistent across multiple biological replicates (Fig S3G and H).

Because these data support the notion that collagen differently impacts migratory phenotypes across cell lines from Black and White women (Fig 4A), we next directly tested invasion potential across the full panel of patient-derived breast cancer cell lines. We first conducted transwell assays with or without collagenous matrigel. We found that both cell lines from Black women are significantly less able to migrate across the transwell compared to those from White women in assays without collagenous matrigel coating (average number of migrating cells/field, White: 48, Black: 12, p= 3.51 x 10^-7^) (Fig 4B). However, precoating transwells with collagenous matrigel largely abrogates these differences in cell invasion between cell lines from Black and White women (average number of invading cells/field White: 13, Black: 9, p= 0.013) (Fig 4C). In fact, the average rate of invasion (invasion in collagenous matrigel compared to migration without a collagen matrix) is significantly higher for cell lines from Black women (White: 0.2779, Black: 0.7872, p= 0.0005) (Fig 4D)

**Figure 4.**
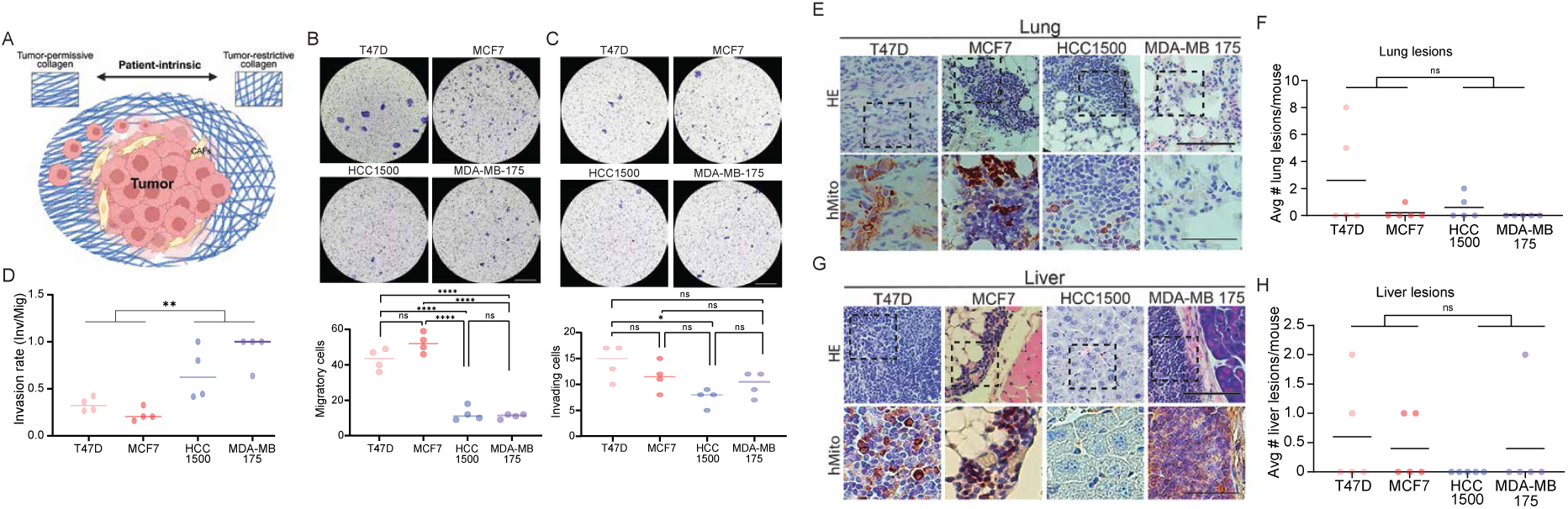
High collagen density and radial organization promotes metastasis of breast cancer cells derived from Black patients. (A) Schematic modeling the effects of collagen deposition and alignment on tumor cell invasion potential. (B-C) Representative images of transwell (B) migration and (C) invasion assays across HR+ breast cancer cell lines from Black and White women, with accompanying quantification represented as bar graphs. (D) Bar graphs depicting invasion rate for transwell invasion assays relative to migration. (E-H) Representative images from H/E (top row) and IHC for human mitochondria (bottom row) of lung (E) and liver (G) micrometastases from xenografts of indicated cell lines with dot plots representing accompanying quantification for lung (F) and liver (H) micrometastases. Scale bars are 290um (B-C) and 140um (E and G). Student’s t-test was used to determine p-values (B-D, F&H), and one-way ANOVA was used with Tukey’s test for multiple comparisons to determine adjusted p-values between groups (B-C). Abbreviations: a.u. (arbitrary units), ctDNA (circulating tumor DNA), HE (Hematoxylin and Eosin), hMito (anti-human mitochondria), um (micron), ns (not significant).

Since we have one experimental model representing collagen organization comparable to tumors from Black breast cancer patients (HCC1500), and one model that does not intrinsically recapitulate this organization (MDA-MB-175) *in vivo*, we next tested whether collagen organization alters metastatic potential in mice. To test the ability of cells to intravasate from the primary tumor into circulation, we assessed ctDNA levels in nude mice xenografted with the panel of cell lines used above, at tumor harvest 12-weeks post xenografting or when tumors had reached 100 mm^3^ in diameter (Fig S4A). In support of our previous observations that high collagen microenvironments promote invasion of Black ER+ breast cancer cells, the MDA-MB-175 xenografted cells, which lack the ability to create a dense, radially-aligned collagen microenvironment, and are unable to form palpable tumors, largely do not have detectable ctDNA levels in their circulation (Fig S4B-C). On the other hand, HCC1500 cells which do recreate a dense and radially organized collagen microenvironment and generate measurable tumors *in vivo*, produce ctDNA levels that are comparable to that from either MCF7 or T47D xenografted tumors, mirroring our *in silico* and *in vitro* results (Fig S4B-C). While we did not identify macrometastases in any of the mice, we were able to detect micrometastases in the lungs (Fig 4E) and livers (Fig 4G) in 40% of the mice (Fig S4D). As with the ctDNA analyses, only one mouse xenografted with the MDA-MB-175 cells shows any evidence of micrometastasis, while HCC, MCF7 and T47D tumors had highly comparable frequencies of micrometastasis incidence (p=0.167 lung, p=0.586 liver) (Fig 4F&H). Of note, the two cell lines from Black women have potential distant site specificity for metastasis, with HCC1500 only demonstrating lung micrometastases, while MDA-MB-175 only metastasize to the liver, unlike the two White lines which metastasize to both sites (Fig S4D). Together, these data suggest that self-reported race is associated with altered tumoral collagen deposition and architecture that likely impacts metastatic potential.

### Patient poverty burden associates with prognostically unfavorable tumor immune landscapes

Because tumoral collagen is an important modulator of immune regulation (Fig 5A) and both pathways (collagen and immune regulation) were systematically identified to be differentially regulated in tumors from Black women by pathway grouping analyses, we next conducted *in silico* CIBERSORTx analysis to agnostically compare predicted immune landscapes from bulk RNAseq of Black and White patient tumors in the VA dataset. Principally, we found signatures reflecting high CD4+ T-helper cell recruitment and depletion of cytotoxic CD8+ T-cell populations in HR+ tumors from Black patients, which is consistent with prior reports^16,17^ (Fig 5B). Predicted populations of B-cells are also higher in HR+ tumors from Black women in this dataset (Fig 5B), also concordant with prior reports^16,17^. We validated these findings in an independent RNA sequencing dataset from the Carolina Breast Cancer Study (CBCS)^6^ with similar corresponding clinical annotations as contained within the VA dataset (Fig 5B, Fig S5A).

**Figure 5.**
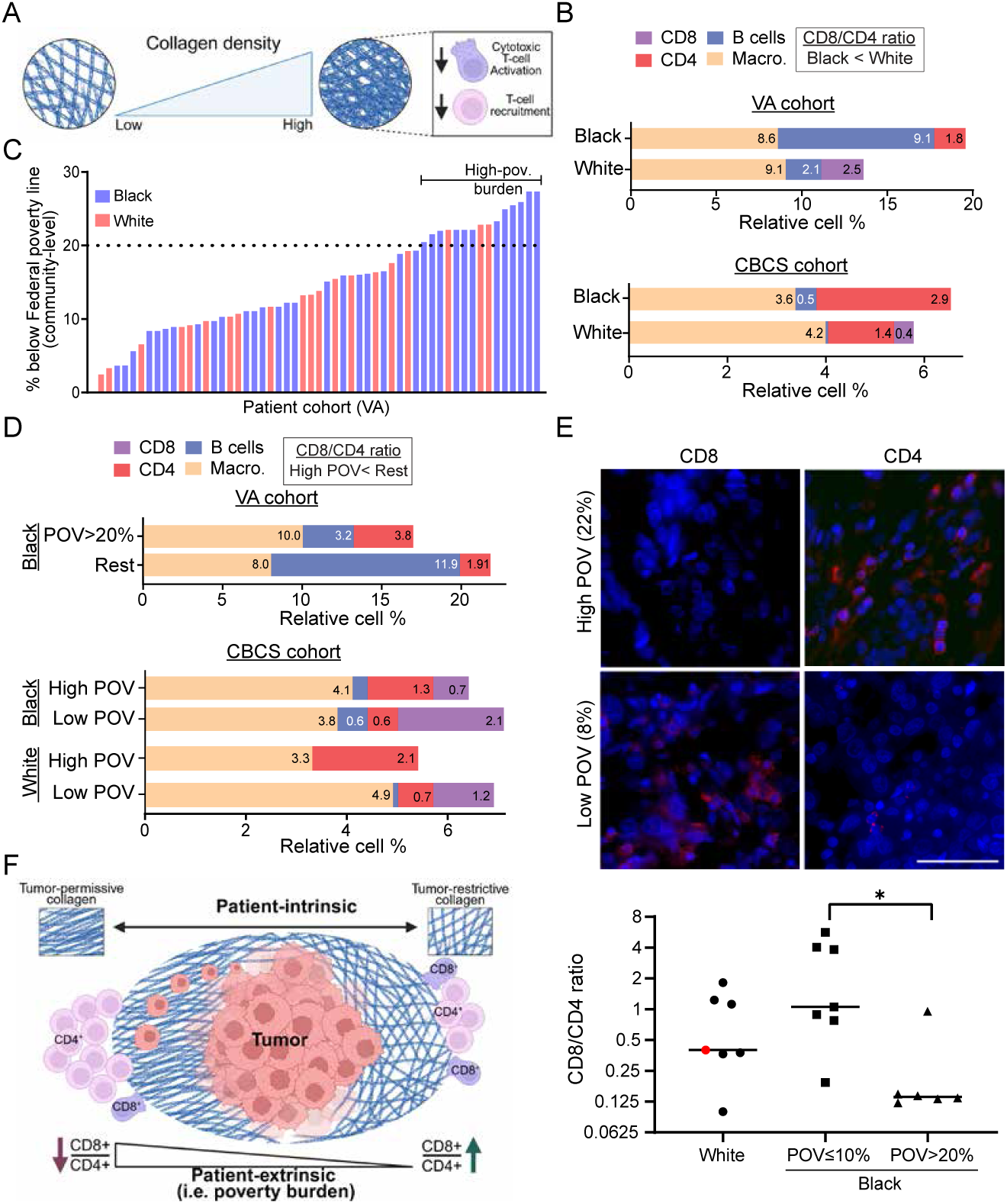
Patient poverty burden associates with an exhausted T cell signature in breast tumors. (A) Schematic modeling known effects mediated by increasing tumoral collagen density on T-cells. (B&D) CIBERSORTx immune cell proportion predictions for major cell groups including CD4, CD8, B-cells, and Macrophages for tumors from Black or White Women (B), and across patient poverty burden indices (D) in indicated datasets. (C) Waterfall plot for patient poverty burden at the community-level by geolocation in the exploratory VA dataset, where high-poverty burden is demarcated by a horizontal dotted line at 20%. (E) Representative images for CD4 and CD8 immunofluorescence of two patient tumors from high- and low-poverty burden in the VA dataset with accompanying quantifications below. (F) Schematic model encompassing tumoral collagen density overlaid with the relationships between patient poverty burden and T-cell recruitment. Scale bar is 40um (E). Student’s t-test was used to determine p-value (E). Statistically significant p values are graphically presented as asterisks, with one asterisk (p<0.05). Abbreviations: CD8 (cluster of differentiation 8), CD4 (cluster of differentiation 4), Macro. (macrophages), VA-cohort (Veterans Affairs cohort), CBCS-cohort (Carolina Breast Cancer Study cohort), POV (poverty burden).

We next investigated the relative contribution of poverty burden to the immune signature observable in HR+ tumors from Black women as poverty burden is reported to trigger chronic stress-mediated immunomodulation^28–30^. In our VA discovery cohort, ∼32% (13/41) of Black women lived in high-poverty burden communities, compared to ∼13% (3/23) of White women. This represents a disproportionate burden that is representative of the larger US population^31,32^, although not statistically significant likely due to sample size limitations (Fig 5C). Poverty burden analyses identified that the exhausted T-cell signature hitherto believed to be race/ancestry-directed, in fact associates with high poverty burden in both Black and White patients in both datasets analyzed (Fig 5D, S5B-C). Of note, enrichment of a B-cell signature frequently observed in Black cancer patients, or those of African ancestry, appears to also be modulated by poverty burden, with enrichment observed in both Black and White patient tumors in low poverty-burden groups (Fig 5D). To validate this observation *in situ*, we performed immunofluorescence on a subset of patient tumors in the VA dataset with appreciable amount of tissue remaining (7 White, and 8 Black) (Fig 5E). This protein-level assay validated the T-cell exhaustion signature (low ratios of CD8+/CD4+ T-cells) as enriched in tumors from women facing high-poverty burden relative to those in low-poverty burden neighborhoods (p=0.012), consistent with CIBERSORTx predictions. Together, these data suggest a potential confluence of stromal collagen alignment and poor-prognostic immune signatures in Black breast cancer patients faced with high-poverty burden which may contribute to the poor outcomes historically associated with this patient subset (Fig 5F).

To identify potential molecular drivers of the association between poverty burden and an exhausted T-cell landscape, we conducted linear regression analyses of the gene candidates identified as being significantly differently expressed between Black and White women in multiple datasets (Fig 1E) and poverty burden. Of the 10 validated candidates, only gene expression of the immunomodulatory regulator *STAT4* inversely correlates with poverty burden in this patient cohort (R^2^= 0.126, p= 0.025) (Fig S6). The correlation appears more robust in tumors from Black women (R^2^= 0.29, p= 0.002) than in those from White women (R^2^= 0.12, p= 0.28) although this may be explained by the wider range of poverty burden faced by Black women compared to White women (Fig 6A). This relationship also validates in the CBCS cohort; HR+ tumors from patients who are below the federal poverty line are significantly more likely to have lower levels of STAT4 expression than those from patients above the federal poverty line irrespective of race (p=0.04) (Fig 6B). Single cell data shows that *STAT4*, which is an immunomodulatory gene required for CD4+ Th1 cell differentiation, is primarily expressed in CD4+ regulatory-T cells, and to a lower extent CD8+ T-cells^33^ (Fig 6C). Analysis of activated pSTAT4 protein levels using immunohistochemistry of patient tumors from the VA cohort similarly identifies enrichment for pSTAT4 positivity in patients with low poverty burden relative to those with high burden (p=0.036) (Fig 6D). Overall, these data suggest a potential role for immunoregulatory genes like STAT4 as molecular drivers modified by lived experiences such as high poverty burden to mediate poor prognostic immune signatures independent of race. In support of this hypothesis, low STAT4 expression significantly associates with poor disease-free survival in HR+/HER2-breast cancer patients in both METABRIC (HR=1.55; p=0.01) and TCGA (HR=3.72; p=0.03) datasets (Fig 6E), in accordance with the literature^34^. Additionally, in the TCGA dataset, stratification of patients by race indicates that both Black (HR=6.4; p=0.002) and White (HR=2.7; p=0.04) HR+ breast cancer patients with low levels of STAT4 have worse survival outcomes than those with higher levels (Fig 6F). Taken together, these data suggest that patients with low poverty burden appear to maintain a functional STAT4 signaling axis, which is critical for promoting a robust Th1-mediated anti-tumor immune response, while patients facing high poverty burden downregulate this axis contributing to worse outcome (Fig 6G).

**Figure 6.**
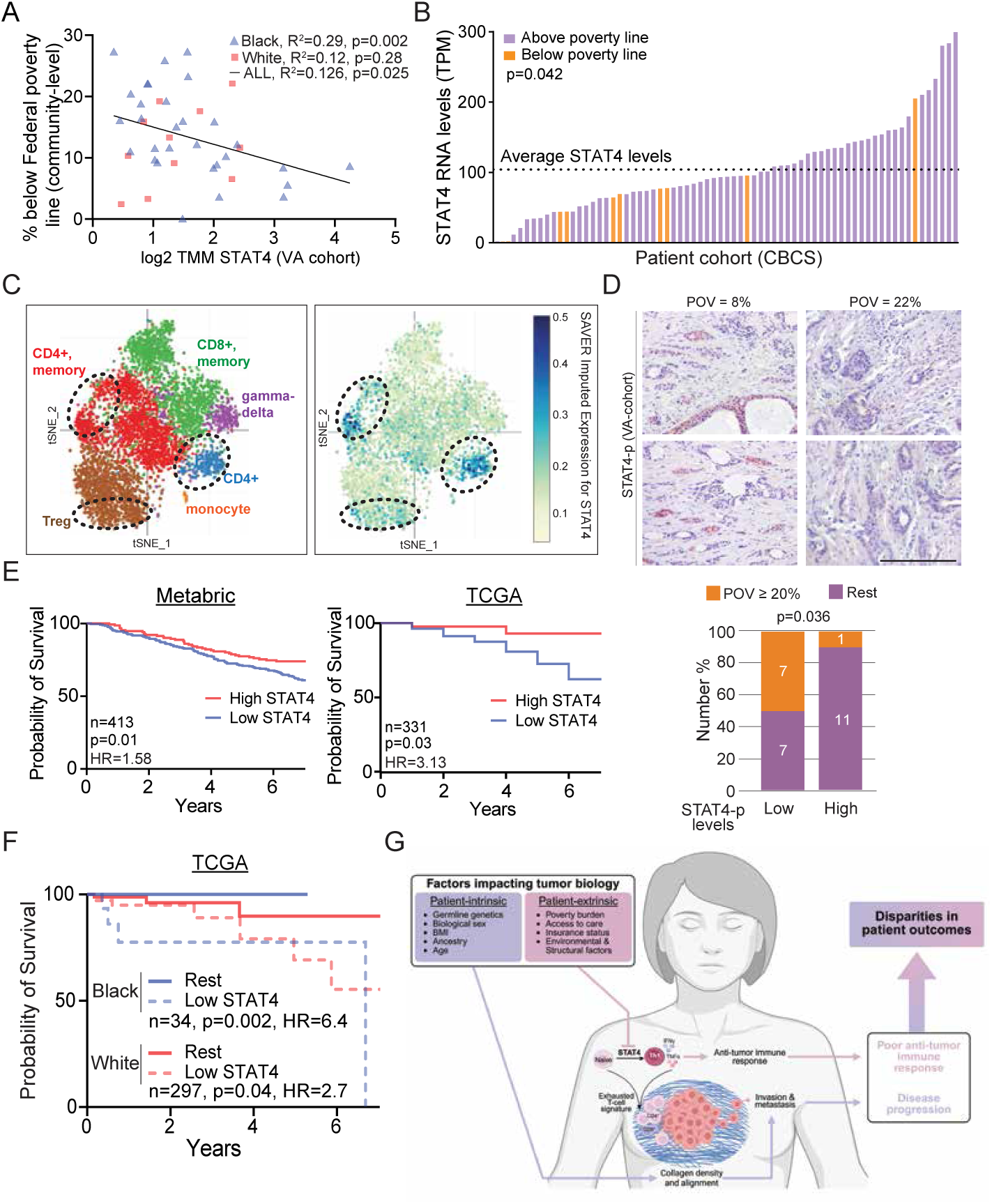
STAT4 as a molecular liaison between poverty burden and poor prognostic immune landscapes in breast cancer patients. (A) Linear regression plot depicting the relationship between STAT4 expression and patient poverty burden in the VA dataset. (B) STAT4 expression relative to patient poverty burden in the CBCS validation cohort, where the horizontal dotted line indicates average STAT4 expression. (C) tSNE plots from the Single Cell Portal for STAT4 expression depicting highest expression levels in immune T-cell subsets. (D) Representative immunohistochemistry images for phospho-STAT4 protein expression in patient tumor sections from the VA-dataset across two separate patient tumors from high- and low-poverty burden with accompanying quantification represented as stacked columns. (E-F) Kaplan Meier survival curves comparing patient tumors with high and low STAT4 RNA expression for women with estrogen receptor positive/human epidermal growth factor receptor 2 negative breast cancer from Metabric (E, left) and TCGA (E, right & F), with (F) or without race stratification (E). (G) Working model schematic highlighting the interplay between patient-intrinsic and -extrinsic factors in the modulation of tumor biology that contributes to poor patient outcomes in Black breast cancer patients. Scale bar is 290um (D). Linear regression coefficients were used to determine p-values using a T-test (A). Fisher’s exact test was used for determining the p-values in (B and D). Survival analysis was performed using the Mantel-Haenszel method to determine hazard ratios (HR) and a log-rank Mantel-Cox test was used to determine p-values (E-F). Abbreviations: TMM (Trimmed Mean of M values), TPM (Transcripts per million), VA-cohort (Veterans Affairs cohort), CBCS-cohort (Carolina Breast Cancer Study cohort), CD8 (cluster of differentiation 8), CD4 (cluster of differentiation 4), Treg (Regulatory T cells), POV (poverty burden), HR (hazard ratio), n (number).

## Discussion

### The importance of integrating life experiences in molecular analyses of patient tumor datasets

In the era of precision medicine, a lack of tumor datasets that are racially, ethnically, and socioeconomically diverse remains a significant barrier to understanding cancer biology across patient demographics. There is also a lack of publicly available molecular and ‘omics tumor datasets with comprehensive corresponding patient annotations that include structural factors like poverty burden, insurance status, access to care, and educational attainment. When structurally annotated datasets exist, they typically do not contain corresponding somatic, molecular transcriptomic, or proteomic data. And yet, there is growing evidence to suggest that integration of social and structural drivers-related data with somatic transcriptomic and proteomic data is critical for implementing precision oncology in the clinic^35,36^.

The data we present here sheds new light on how patient characteristics such as race drive tumor cell-intrinsic somatic signaling to promote cancer relevant phenotypes such as ECM organization in the microenvironment, which significantly impacts metastatic potential. In parallel, we show that life experiences such as high poverty burden can drive poor prognostic immune landscapes with exhausted T-cell signatures to inhibit anti-tumor immune response and contribute to poor prognosis for the patient. To our knowledge, this report is among the first to deconvolute the impact of life factors such as poverty burden from that of ancestry by characterizing the interplay of these complex factors through the lens of somatic molecular biology and experimental model systems. The importance of such findings is exemplified by our discovery that the exhausted T-cell signatures previously attributed to ancestry is in fact driven by life factors such as high poverty burden which are often deeply entwined with race. While poverty stressors are complex sociological problems that are currently therapeutically unaddressable, perhaps identifying poverty-associated biomarkers could eventually guide stratification of patients for clinical decisions surrounding immunotherapy. Undoubtedly, these data provide proof of concept for an imperative and urgent need to generate more fully annotated datasets that represent real world patient demographics and life histories, in combination with molecular ‘omics, if we are to achieve the goal of precision oncology for all.

Deconvoluting self-reported race from genetic ancestry poses an additional hurdle rapidly gaining recognition in genomics and associated fields. For example, the African diaspora is the most genetically diverse demographic in the world^37,38^ and yet is frequently monolithically annotated in racial constructs as Black. Despite considerable overlap, self-reported race and genetic ancestry are not synonymous, but can each provide different insight into tumor biology relevant to patient outcome. For instance, in the VA dataset described here, we identify two subsets of Black patients (patients above and below median line, Fig 2B) with differing levels of tumoral collagen in which one group is primarily driving the elevated differences compared to White patients (FC= 3.77, p=0.002). This may well be attributable to differences in ancestral genetics amongst women who self-identify as Black, although this dataset does not have the genomic resolution to interrogate this hypothesis. However, using self-identified race as a metric can provide additional clinically relevant insight into confounding structural and cultural factors that may be missed with a myopic focus on ancestry alone. Future investigations incorporating both ancestral genetics and self-reported race in larger patient cohorts are certainly warranted to test this, and other such hypotheses.

Perhaps the most striking finding presented here is the discovery that poverty aligns with a poor prognostic exhausted T-cell signature. Several important previous studies have identified the exhausted T-cell landscape detectable in breast tumors of Black women as a race-aligned characteristic ^16,20^. However, in the US, Black people are nearly twice as likely to live in poverty compared to White counterparts^9,32^. Therefore, should poverty burden contribute to this immune phenotype, then Black women would indeed be at increased risk, but so would many other patients irrespective of racial or ethnic identify. While virtually unstudied in cancer biology, this discovery of poverty-influenced immune signatures should not be surprising. Several studies across disciplines have reported an impact for patient-extrinsic factors of socioeconomic status on immune responses^39^, signaling^39^, and differentiation^40^. For example, low socioeconomic status (education, income, neighborhood deprivation index, etc.) is associated with elevated levels of pro-inflammatory cytokines and additional molecular drivers underlying chronic inflammation^41,42^, and defective CD4 T-cell differentiation^39^. Conversely, high socioeconomic status is reportedly associated with increased inflammatory immune signaling through upregulation of interferon gamma^40^, a cytokine largely produced by Th1 CD4+ T-cells, which is dependent upon a functioning IL12/STAT4 signaling axis. Development of experimental model systems and biomarkers that account for life experiences is a critical next step for precision oncology.

Importantly, immune landscapes are also impacted by the ECM composition within surrounding tumor microenvironments and actively influence ECM remodeling. Therefore, patient-intrinsic differences in the ECM of tumors across different patient demographics pose an additional layer of complexity that undoubtedly impacts tumor behavior and patient outcomes. This is historically observed across many investigations reporting that breast tissue from Black women is more often categorized as having increased fibrosity and density^43^, which are negative prognostic factors correlating with more aggressive tumor behavior, and contributing to less effective early detection screening by mammography^44^. In the context of the rapid uptake of generative Artificial Intelligence models to read mammogram reports, better understanding of race-related collagen characteristics is of critical importance to address bias in models^45^.

### Race-aligned tumoral collagen organization is critical for cancer cell invasive potential in Black women

Collagen is a primary ECM component contributing to increased breast density^46^ and tissue fibrosity as fibers become increasingly aligned^47^. Similarly to the results reported here of increased radial organization of collagen fibers in tumors from Black HR+ breast cancer patients, recent work also identified high collagen density and fiber alignment in triple negative breast tumors from Black women^48,49^. Here, we further report, and for the first time, the ability of tumor cells derived from Black women to direct collagen density and organization through cell-intrinsic signaling. Indeed, a HR+ breast cancer cell line derived from a Black woman, which is unable to direct these collagen changes in the breast microenvironment is incapable of generating defined, palpable tumors in a xenograft model. These results suggest that tumor cells in Black women may be habituated to, and require, dense, radially aligned collagen environments to achieve growth and progression. This has important implications for commonly used cancer research experimental model systems that almost completely ignore collagen, and therefore, may be ill-equipped to support the growth of cancer cells from Black women.

Additionally, *in silico* predictions based on patient data, as well as *in vitro* and *in vivo* experiments suggest that the collagen environment housing tumors in Black women is critical for promoting metastatic progression. Black breast cancer is often characterized as more metastatic, either because Black women present with more advanced cancer or because of the more aggressive proliferation reported in Black breast cancer patient tumors^17,18,21^. Here, for the first time, we present evidence supporting no intrinsic difference in invasive potential of HR+ breast cancer cells growing in Black and White women. Rather, these cells have evolved to identify methods of invasion and metastatic growth suited to the ECM that they grow within. Collagen density and organization could certainly be utilized as prognostic biomarkers to be tailored for different patient demographics in efforts to better predict metastatic potential. Further mechanistic investigations into the cell-intrinsic signaling that drives these collagen patterns and their implications for drug delivery and efficacy are warranted.

While the results presented here provide several new and striking insights into the somatic molecular biology underlying poor outcomes for Black breast cancer patients, there are several caveats to be considered. While the dataset generated here is uniquely well annotated for many patient characteristics and largely controlled for access to health care and education levels, the sample size is relatively low. Therefore, this study is only powered to detect the most readily observable molecular changes, which are likely to represent fundamental key drivers of outcome disparities. There are undoubtedly many other features unique to breast cancer in patients from different demographics within either the Black or White communities that this dataset is not sufficiently powered to identify. Additionally, in several instances, the experimental approaches to test causality were restricted by available resources. Currently, cancer research techniques and model systems are tuned to foster the growth of cancer cells derived from White breast cancer patients and therefore are ill-suited to address hypotheses derived from data from Black women. For instance, collagen alignment appears to differently impact cancer cell migration phenotypes between cancer cell line models from Black and White women. While we were able to identify breast cancer cell lines from Black women that could replicate some of the phenotypes observed in patients both *in vitro* and *in vivo*, these experimental approaches need substantial refinement. Moreover, although we were able to identify quantify baseline invasion rates for cell lines *in vitro* using matrigel, a comparison between matrigel and no matrigel likely introduces several confounding factors other than collagen, which must be better controlled for in order to comprehensively investigate the impact of collagen on the invasion capacity of cancer cell lines *in vitro*. Understanding the exact ECM changes and engineering tissue culture systems that incorporate these findings to better replicate the microenvironment of Black breast cancer is a call to arms for all cancer researchers. By integrating social and structural drivers of health with underlying molecular impacts, we may begin to be able to advance better biomarkers and treatment plans tailored to the patient. Achieving truly inclusive precision oncology will likely require better integration of strategies to expand diverse patient datasets, develop representative experimental models, and generate hypotheses that reflect the full spectrum of cancer biology and human experiences.

## Materials and Methods

### Datasets

#### Tumor datasets

The primary tumor dataset presented in this study was obtained from the Veterans Affairs office in Durham, North Carolina. All tumors were molecularly characterized as hormone receptor positive (ER and/or PR positive) comprising a total of 64 archival tumors (Black n=41, White n=23). Corresponding patient annotations were obtained and included both clinicopathological characteristics and SDOH (n=60). Validation datasets included in this study are sourced from the publicly available TCGA (downloaded August 2023), Metabric (downloaded August 2023), Ellsworth (GEO78958), and the Carolina Breast Cancer Study (CBCS). All validation datasets were treatment-naïve. RNA sequencing, corresponding clinicopathological annotations and patient outcomes for TCGA and METABRIC were downloaded from cBioPortal (downloaded June 2023). The CBCS tumor dataset was obtained from collaboration with Dr. Melissa Troester at University of North Carolina, Chapel Hill (downloaded February 2024).

#### VA cohort clinicopathological and socioeconomic annotations

All patients were diagnosed and treated within the VA healthcare systems surrounding Durham, North Carolina between 1997 and 2020 with continued surveillance monitoring disease recurrence, metastasis and death. BI-RADS scoring system was used for initial pathological characterization. Subsequently, either TNM tumor stages or tumor grades were reported for each patient. Histopathological subtyping was performed by IHC for HER2, ER, and PR. Each patient profile contained additional socioeconomic indicators. Body mass indices were reported for each individual. Additionally, community-level metrics were reported for poverty burden, household income, %-employment, and %-high school graduate. These community level indicators were determined by geocodes for each patient. For example, a 20% poverty burden indicates that a patient resides in a high poverty burden community where 20% of people are below the Federal poverty line, etc.

#### Bulk RNA sequencing and gene set enrichment analyses from patient tumors in the VA cohort

56 archival FFPE patient tumors (Black=36, White=20) yielded high-quality RNA following standard extraction methods and were processed for bulk RNA sequencing using next-generation sequencing services through Novogene. For RNA analysis, p values were obtained by comparing each gene between tumors from Black and White women using the two-tailed Wilcoxon Rank Sum test within each dataset analyzed. Differentially expressed genes for the VA dataset were grouped into moderate fold-change (FC 1-1.5) and high fold-change (FC>1.5) and multiple orthogonal gene set enrichment analyses were performed. Reactome Pathway Analysis web tool (https://reactome.org) was utilized to identify differentially regulated pathways mapping to genes with moderate and high fold-change expression. These were validated in both Ingenuity Pathway Analysis (IPA) and Metascape pathway grouping analysis. Differentially expressed gene candidates were cross-validated in two independent datasets (TCGA and Ellsworth) to refine RNA candidates with standard cut-offs of mean +/- 1 standard deviation (SD) to validate ‘high’ vs ‘low’ expression candidates.

#### Bulk RNA sequencing and gene set enrichment analyses from xenograft tumors

Frozen xenografted tumor samples were submitted to Azenta Life Sciences for RNA extraction, library preparation, sequencing, and alignment to the human reference genome (GRCh38) using their standard pipelines. Gene-level transcript abundances were normalized to transcripts per million (TPM) values. TPM values were used to identify genes with fold changes greater than 1, which were subsequently submitted to the Reactome Pathway Analysis web tool (https://reactome.org) for enrichment analysis. Raw count data were used for downstream differential expression analyses in RStudio (v2025.05.1+513). Principal component analysis (PCA) was performed using the variance stabilizing transformation (vst function, DESeq2 v1.44.0) to evaluate global transcriptional variation between samples. Differential expression analysis was carried out using DESeq2 with default normalization and dispersion estimation parameters. Differentially expressed genes were defined as those with an adjusted p-value<0.05 (Benjamini–Hochberg correction). Heatmaps of the top 100 most significantly differentially expressed genes were generated using the pheatmap package (v1.0.13).

#### Carolina Breast Cancer Study RNA sequencing validation dataset

CBCS3 tumor RNA was extracted from 2 1-mm formalin fixed, paraffin-embedded cores using the Qiagen RNeasy FFPE Kit (Cat# 73504)^50^. Sequencing libraries for 79 samples were prepared from 1.0 µg of total RNA at the UNC Translational Genomics Laboratory (TGL) using a Hamilton STAR Automated Liquid-Handling Platform and the TruSeq Stranded Total RNA Library Prep Gold Kit following manufacturer protocol. Library quality was measured using a TapeStation 4200, while RNA quantity was measured using a Qubit 3.0 fluorometer. Total RNA libraries were pooled in equal molar ratios and sequenced on an Illumina NovaSeq6000 sequencer with 2x50 bp paired end reads to an average sequencing depth of ∼125 million clusters; these were used to generate raw sequencing read files (FASTQs).

#### Survival plots

For Kaplan–Meier survival curves, all patients with associated survival data in the VA dataset were used, and HR+/HER2- patient subsets were used in the TCGA and METABRIC validation datasets. Outcome measures used were progression-free survival (integrating recurrence, metastasis, disease-specific). Outcome measures were selected to maximize sample size. Only samples with survival metadata were included in the analysis. Log rank test calculated p values. Standard cut-offs for STAT4 expression as a differentially expressed candidate were mean +/- 1 SD for TCGA and METABRIC datasets. Survival analysis for STAT4 expression stratified by self-reported race in the TCGA dataset was performed using the expression cut-off of mean - 1 SD (low) compared to ‘Rest’.

### Histology and Immunostaining

#### Collagen staining and quantifications

Collagen levels were assessed via Masson’s Trichrome in a subset of patient tumors from the VA dataset containing sufficient tumor tissue following bulk RNA sequencing. Sections were imaged using the ECHO Revolve microscope. Representative images were taken at 10x magnification and images for second harmonic generation (SHG) were taken at 40x. Manual collagen quantifications were conducted via FiJi after colour_deconvolution to separate collagen content from cells. Collagen intensities were grouped into high and low-intensity collagen stains to allow for accurate collagen quantifications after setting threshold gates to ensure maximum collagen detection across all samples. Within each bin, collagen intensity was thresholded consistently across tissue samples. Collagen angles surrounding tumoral ducts were measured using the ‘angle tool’ in FiJi by selecting 5 representative collagen fibers relative to ductal cell intersection (Black n=9, White n=7).

Second harmonic generation was performed as an agnostic and orthogonal validation to the manual quantifications. For this, trichrome-stained images from White and Black patient tumors were analyzed to measure collagen area, density, and fiber alignment on NIH Image J. The colour_deconvolution plug-in in ImageJ was used to separate the respective color channels: red, blue and green (RGB). To measure collagen area and density, blue-channel images were converted to 8-bit grayscale format to enhance contrast. The Threshold tool was used to select and highlight collagen regions, adjusting the threshold levels to accurately differentiate collagen from non-collagen areas. Collagen area and integrated density were measured using ImageJs measurement tools. To analyze collagen fiber orientation, the blue channel images were isolated in ImageJ. A single-pixel-line filter was applied at sequential rotational intervals (0°, 1°, 2°, etc.), and the brightness of the remaining pixels was summed^51^. The summed brightness at each angle was computed into a fraction, with the total of all fractions equaling 1. Angles were then grouped into increments of 30° by adding the fractions from 1° to 30°, 30° to 60°, and so on, to assess angle distribution trends.

#### Immunohistochemistry and Immunostaining

Immunohistochemistry was performed according to the manufacturer’s instructions. Sections were first deparaffinized, then endogenous peroxidases were quenched using 3% H2O2, and antigen retrieval was done using 1x citric acid buffer. The blocking buffer used was 2% goat serum. Antibodies used were STAT4 (Cell Signaling Technology, catalog no. #4134), and Histone H3 (Cell Signaling Technology, catalog no. #9701). Primary antibodies were left overnight in 4° and followed by anti-rabbit secondary (Vector Laboratories, catalog no. BA-1000) or anti-mouse secondary (Vector Laboratories, catalog no. MKB-2225). Next, sections were incubated in avidin-biotin complex solution (Vector Laboratories, catalog no. PK-6100), stained with peroxidase substrate (Vector Laboratories, catalog no. SK-4800), and counterstained in hematoxylin. Immunofluorescence was performed according to the manufacturer’s instructions and as per previously published protocols^52^. Antibodies used include CD206 (Thermo Fisher Scientific, catalog no. MA5-16871), CD86 (Thermo Fisher Scientific, catalog no. PA5-114995), CD8 (Thermo Fisher Scientific, catalog no. MA5-13473), and CD4 (Abcam, catalog no. ab133616). Images were captured on an Echo Revolve microscope.

### Cell culture

#### 2D migration and invasion assays

Transwell migration assays were performed to assess metastatic potential comparing HR+ cell lines derived from Black (HCC1500 and MDA-MB-175) and White (MCF7 and T47D) patients. For transwell migration and invasion assays, 25,000 cells were plated in 200 μl of media (without fetal bovine serum (FBS) supplementation) in 8uM transwell inserts (Falcon, catalog no. 353182). For invasion assay, inserts were pre-coated with Matrigel (Corning, catalog no. 356234) in 1:3 dilution with media without FBS. Inserts were placed on a 12-well plate with 750 μl of standard cell culture growth media and growth for 24 hours. Then, transwells were fixed and stained using HEMA3 staining kit (Thermo Fisher Scientific catalog no. 22-122911). Inserts were dried, and pictures were taken using the Echo Revolve microscope at 10x magnification. Invading cells were manually counted using FiJi.

### *In vivo* experiments

#### Xenograft tumor growth assays

Four-week-old nude mice (from JAX) were used for all tumor xenografts (5 mice per group, 20 mice total). Mice were given estradiol-supplemented sterile deionized water at a concentration of 8 μg/ml starting one week prior to tumor injections and throughout the experimental duration. At five weeks of age, each mouse was injected bilaterally in #4 mammary fat-pads with 3 million cells per injection resuspended in 200uL of a 1:1 mixture of Matrigel (Corning catalog no. 356234) and standard growth media. Mice were palpated weekly until tumors were palpable. Once a tumor was palpable, palpations were done twice a week for 12 weeks or until tumors reached 100 mm in diameter. Mammary fat pads were harvested for all mice without a palpable tumor for HE histological assessments. Tumors and fat pads were fixed in 4% paraformaldehyde and paraffin embedded prior to histological assessment and immunostaining.

### *In silico* immunotyping

Following recommended usage guidelines, immune landscape predictions for major cell components were determined using CIBERSORTx with 100 permutations (1000 permutations showed no change in proportionality), and quantile normalization disabled. We used the LM22 as the primary reference matrix for all major immune cell populations grouped to include CD4-positive, CD8-positive, B cells, Natural killer cells, and Macrophages. We utilized the Single Cell Portal (https://singlecell.broadinstitute.org) for the analysis and visualization of STAT4-expressing cells at the single-cell level, revealing primary expression in T-cells. Datasets were accessed and processed using the publicly available tools and resources located on the portal.

### 3D invasion modeling

The major aim of this computational model is to bridge transcriptomic data with cell migration behaviors through a three-module pipeline, described in the following sections. This framework enables the investigation of how transcriptomic states—particularly those related to cytoskeletal and mechanotransduction machinery—dictate cell mechanics and invasive potential at the patient level.

Transcriptomic data from non-Hispanic Black and White women with breast cancer, from the discovery VA-cohort, have been incorporated into the model to predict potential migration trajectories under varying extracellular matrix (ECM) conditions, specifically across different collagen fiber densities and alignments. This approach offers a mechanistic understanding of how gene expression differences may contribute to phenotypic diversity in cell motility and invasiveness within racially diverse patient populations.

Module 1:

The first module serves as a critical interface between the transcriptomic and proteomic scales by discretizing gene expression levels and filtration of gene groups. It is designed to work with both bulk and single-cell RNA sequencing datasets and requires at least two distinct cellular states for comparative analysis. Initially, cytoskeleton-related genes are identified from the transcriptomic data and analyzed to compute both p-values and log₂ fold changes between the states. These two metrics are then used to categorize gene expression levels as high, medium, or low, based on intermediate reference states. In addition to the two observed cell states, the signaling simulation incorporates intermediate, hypothetical states in which genes are expressed at medium levels and exhibit no significant differences between groups. Genes that pass the filtering thresholds for p-value and log₂ fold change are included in gene sets that show either high expression in group II and low in group I, or vice versa. Significant genes with an absolute log₂ fold change greater than 1.2 are classified as highly expressed and assigned a value of 1. Their corresponding expression in the opposing state is set to 0.0001, while reference genes are assigned a value of 0.01. These assigned values serve as fixed initial states for the cytoskeletal signaling network in Module 2. This bidirectional analysis enables a precise, state-specific representation of cytoskeletal signaling, forming the foundational input for phenotype-dependent modeling of cell behavior.

Module 2:

Module 2 executes the intracellular signaling simulation using our previously established computational framework, which integrates cytoskeletal signaling and upstream regulatory networks^53^. The cytoskeletal dynamics model captures emergent biophysical behaviors by estimating key parameters such as actomyosin contractility, F-actin polymerization kinetics, integrin binding and unbinding rates, crosslinker-mediated F-actin bundling, cell polarization, and effective cellular stiffness each of which plays a critical role in characterizing invasive phenotypes of cancer cells. These parameters are governed by extended Boolean differential equations, a formulation capable of handling large-scale signaling networks while incorporating the mechano-chemical properties of cellular functional groups. The discretized gene expression values derived from Module 1 are incorporated into this system through logical conjunctions (AND operations), forming a set of discrete, time-dependent ordinary differential equations (ODEs). This integration enables dynamic modeling of transcriptional regulation and its impact on the cell’s physical interactions with its environment.

Module 3:

Following Module 2, Module 3 introduces a physical model that connects cytoskeletal parameters to corresponding 3D biophysical attributes and uses these to then predict ECM characteristics dependent cell migration and invasion. The cytoskeletal parameters obtained from the signaling simulation are scaled to their physical counterparts using literature-derived ratios. Each attribute is constrained by assumed biological minimum and maximum limits to ensure physiological relevance.

This model builds upon our previous single-cell simulation frameworks^54^. Specifically, the 3D single-cell model is designed to study path persistence—how consistently a cell maintains its migration direction within a 3D fibrous extracellular matrix (ECM).

In this computational model, the extracellular matrix (ECM) is represented as a dynamically generated fibrous network. Rather than predefining the entire matrix structure, fibers and fiber binding sites are stochastically generated in real-time as the cell migrates, allowing for memory-efficient simulations of large-scale microenvironments. Each fiber is characterized by defined directionality and spacing, with its physical properties sampled from probability distributions that reflect specific ECM density and fiber alignment parameters. The matrix can be tuned to replicate diverse microenvironmental conditions, such as isotropic (random) networks or anisotropic (aligned) architectures.

The Alignment Index (AI) quantifies the degree of fiber alignment relative to a reference axis, defined here as the vector v_ref_ = [1, 0, 0]. When AI = 0.0, fibers are oriented randomly; when AI = 1.0, all fibers are perfectly parallel to the reference vector. For intermediate alignment (AI > 0.0), the angular orientation of each fiber is sampled from a normal distribution, where the standard deviation σ_f is determined by the AI value according to the following relationship (Eq.1)^74^.

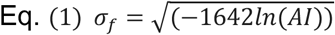

Within the model, three key collagen parameters can be independently tuned: Collagen density, collagen stiffness, and collagen alignment. Collagen density of 1 mg/ml, 2 mg/ml and 4 mg/ml are selected to mimic the low-, medium- and high-density matrices^55^. Collagen stiffness is set to 30,150 and 600 Pa matches physiologically relevant elastic modules based on prior experimental measurements respectively^56,57^. Alignment index is set to AI=0.4, consistent with approximate cell embedded alignment observed in experimental systems^58^. Overall, the cell is modeled as a point agent migrating in continuous 3D space (off-lattice). It forms protrusions to probe nearby ECM fibers and selects one as a target for forward movement. Migration direction is influenced by internal polarity, allowing the cell to exhibit persistent movement by favoring directions aligned with its prior trajectory.

### Reconstitution of 3D Collagen Matrices with 30° Inclined Surface

To reconstruct 3D collagen matrices to have aligned and unaligned collagen fibers, we adapted the protocol described in Sapudom et al^59^. Collagen solutions were prepared in 35mm cell culture dishes (83.3900, Sarstedt). Using Autodesk’s Fusion360™ software (Autodesk Inc., San Rafael, CA, USA), a surface with an incline of 30° was printed using the Bambu Lab X1 Carbon 3D printer (Bambu Lab, Austin, TX). We added 250uL of 3 mg/mL Bovine Collagen Type I Solution (Advanced BioMatrix, Carlsbad, CA, USA). Dishes were placed on the 30° inclined surface in the incubator for 20 mins at 37°C until the collagen had fully dried. For unaligned surfaces, dishes with collagen were incubated without inclination.

### Digital Holographic imaging

The Holomonitor M4 digital holographic microscope was purchased from Phase Holographic Imaging (PHI, Sweden) and placed directly in an Isotemp 3532 incubator (Thermo Scientific). T47D and HCC1500 were cultured in 35mm cell culture dishes (83.3900, Sarstedt). Cells were counted using the Countess 3 Automated Cell Counter (Thermo Fisher Scientific, Waltham, MA, USA). 5000 cells were cultured for 24 hours prior to enable adhesion and each experiment for the cell lines were repeated three times.

Dishes were covered with HoloLids (PHI, Sweden) to stabilize imaging conditions and placed in the microscope for 24 hours before imaging. After 24 hours, 15 imaging positions were selected per cell line and collagen alignment using the HOLOMONITOR App Suite software. The experiments were repeated three independent times, we analyzed a total of 3255 cells for HCC1500 unaligned collagen, 2840 cells for HCC1500 aligned collagen, 2819 for T47D unaligned collagen, and 2533 for T47D aligned collagen condition.

Images were analyzed on the Holomonitor App Suite software (PHI, Sweden) using the “In-depth Analysis: Cell Morphology” feature to segment and quantify cell morphology differences. The segmentation and quantify feature identified how many cells were imaged as well as the change in their area, volume, eccentricity, and irregularity.

### UMAP and Clustering Analysis of Cell Morphology Data

To further distinguish the morphological heterogeneity between unaligned and aligned collagen conditioned, further computational analysis was performed using Python(v3.11) with the pandas (2.2.2), scikit-learn (1.6.1) and umap-learn (0.5.9.post2) libraries.

To create the scatter plots, all control replicates were combined to form a single reference dataset. The parameters from the combined control dataset were standardized using StandardScaler function from scikit-learn. Using the Uniform Manifold Approximation and Projection (UMAP) algorithm, a two-dimensional embedding was created from the standardized control dataset. Embedding the UMAP algorithm gave definitive phenotype charateristics for all subsequent classifications. The control dataset was the unaligned collagen condition. The parameters used from the Holomonitor data were Cell Area, Cell Volume, Eccentricity, and Irregularity.

Performing K-Means clustering (k=4 clusters) on the resulting two-dimensional UMAP coordinates of the control cells identified cell populations. The center of the cluster and boundaries were defined based on the phenotypic distribution of the control population.

Cells in aligned collagen were compared against the unaligned collagen reference map by standardizing the dataset from all aligned collagen replicates using the same scaler previously fitted to the control unaligned dataset. Using the trained UMAP model, the scaled aligned collagen data was projected into the established UMAP. Finally, each cell from the aligned collagen group was categorized in one of the four defined clusters in the control group using the pre-trained K-Means model. To calculate the total cell population as a percentage, the relative abundance of cells within each cluster was calculated for each individual experimental replicate pair (unaligned vs. aligned). This process was repeated independently for each cell line.

### Statistical analyses

Linear regression analyses were used to evaluate the correlation between patient poverty burden and expression of gene candidates validated to be significantly differently expressed in tumors from Black women compared to White women. The co-efficient of determination (R^2^) was calculated as 1 minus the ratio of residual variance to total variance to compute goodness-of-fit, and p values were calculated by an F test. STAT4 expression in the VA dataset was the only candidate identified as being significantly correlated with poverty burden in the VA dataset (R^2^= 0.123, p=0.025). Categorical poverty burden validation of STAT4 expression correlation in the CBCS dataset was performed using Fisher’s exact test of above vs below the federal poverty line for each patient in the dataset (p=0.042). Categorical tests with sample sizes above 15 (per group) were conducted via chi-squared analysis, sample sizes less than 15 were conducted with Fisher’s exact tests. Survival analysis was performed using the Mantel-Haenszel method to determine hazard ratios (HR) and log-rank Mantel-Cox tests were used to determine p-values. Students’ T-tests were conducted for all continuous data comparisons between two groups. For comparisons of continuous data across 3 or more groups, a one-way ANOVA was used with Tukey’s test for multiple comparisons to determine adjusted p-values between groups, when necessary. All *in vitro* assays including transwell migration, invasion, and 3D spheroids were performed and reported in three biological replicates. Statistically significant p values were graphically presented as asterisks, with one asterisk (p<0.05), two asterisks (p<0.01), three asterisks (p<0.001), and four asterisks (p<0.0001).

