## Supplementary Figure Legends for "The impact of patient biology on racial disparities in breast cancer outcome"

Supplementary figures and legends

Figure S1. Hormone receptor positive breast tumors from Black women differently regulate pathways underlying extracellular matrix and immune signaling compared to those from White women. (A) IPA summary plot for pathways recognized as being significantly differently regulated in tumors from Black women. Pathways in bold are overlapping pathways determined from Metascape grouping analysis. (B) Reacfoam plots depicting differentially regulated pathway clusters between tumors from Black and White women grouped by high- and low-fold change expression. Supports data presented in Fig 1D. Statistically significant p values were graphically presented as asterisks, with one asterisk (p<0.05). Abbreviations: VA (Veterans Affairs), adj. (adjusted), TMM (Trimmed Mean of M values), ECM (extracellular matrix), FC (fold change), and IPA (Ingenuity Pathway Analysis).
Figure S2. Hormone receptor positive breast tumors and cell lines from Black women have differently organized collagen compared to those from White women. (A) Linear regression plot for collagen content as a function of BMI in the VA discovery cohort. Pink squares represent White patients and blue circles represent Black patients. (B) Representative HE images from xenografted tumors present in the mammary fat-pads following tumor harvests. Scale bar is 170um. (C) Tumor growth curves for cell lines xenografted into fat pads of nude mice. (D) Tumor growth rate box-plot for xenografts with a horizontal line indicating median tumor growth rate with upper and lower bounds representing the 75th and 25th percentile, respectively. Whiskers reflect the minimum and maximum tumor rates. (E) Principal Component Analysis for xenografted tumors. (F-G) Summary plot for Reactome pathway grouping analysis of differentially regulated RNAs in HCC1500 vs MCF7 (F), and HCC1500 vs T47D (G) tumor xenografts. Linear regression coefficients were used to determine p-values using a T-test (A) and Student’s t-tests were used to determine p-values (C).  Abbreviations: cyto (cytoplasm), mm (millimeter), ns (not significant), PCA (Principal Component Analysis).
Figure S3. Hormone receptor positive cell lines from Black women require collagenous Matrigel for 3D spheroid formation. (A-B) Quantification of 3D spheroid size for tumor cells from HR+ breast tumor cells from Black or White women grown (A) in the absence of an extracellular matrix (soft-agar), or (B) presence of an extracellular matrix (matrigel). (C-D) Effect-size plots of spheroid size in soft-agar compared to matrigel for cell lines from (C) White and (D) Black women. Each dot represents the average size of at least 20 spheroids from an independent biological replicate for three biological replicates per cell line per condition. (E) Representative holomonitor images for cell lines in aligned or unaligned collagen conditions. (F) Heat-maps for cell features used for UMAP cell clustering including cell area, optical volume, eccentricity and irregularity in aligned collagen conditions relative to unaligned. High eccentricity and irregularity indicate clusters corresponding to migratory populations. For T47D migratory clusters correspond to cluster zero and 2. For HCC1500, migratory clusters correspond to clusters one and three. (G&H) Stacked bar graphs depicting the relative proportions of cells in migratory clusters (shades of pink) or non-migratory clusters (shades of grey) in both aligned and unaligned collagen conditions for all three biological replicates. Scale bars are 100um (E). One-way ANOVA was used with Tukey’s test for multiple comparisons to determine adjusted p-values between groups. Statistically significant p values were graphically presented as asterisks, with one asterisk (p<0.05), two asterisks (p<0.01), and three asterisks (p<0.001). Abbreviations: um (micron), ns (not significant), SA (soft-agar), Mat (matrigel), Col-3 (3 mg/mL collagen 1), UA (unaligned collagen), Al (aligned collagen), A (cell area), V (cell volume), Ir (cell irregularity), Ec (cell eccentricity).
Figure S4. Hormone receptor positive patient-derived cell lines from Black and White women can form micrometastases in the lung and liver. (A) Proportion of mice with detectable ctDNA levels (asterisks) after xenografting of breast cancer cell lines derived from Black (HCC1500 and MDA-MB-175) and White women (MCF7 and T47D) into nude mice. (B-C) Dot-plots representing relative ctDNA levels (B) and ctDNA levels normalized to tumor burden (C) in xenografted mice at tumor harvest. (D) Stacked bar graphs for frequency of micrometastasis detected in xenografted nude mice across cell lines. Abbreviations: a.u. (arbitrary units), ns (not significant), HE (Hematoxylin and Eosin), L (left), R (right), ctDNA (circulating tumor DNA).
Figure S5. Patient poverty burden associates with poor prognostic tumor immune landscapes. CIBERSORTx immune cell proportion predictions for major cell groups including CD4, CD8, B-cells, and Macrophages in patient tumors depicted as summary tables (A&C) and a stacked column graph (B) for indicated groups. Abbreviations: CD8 (cluster of differentiation 8), CD4 (cluster of differentiation 4), Macro. (macrophages), VA (Veterans Affairs cohort), CBCS  (Carolina Breast Cancer Study cohort), POV (poverty burden).
Figure S6. STAT4 correlates with poverty burden for patients with hormone receptor positive breast cancer. Summary table for linear regression analyses of 10 validated candidate genes found to be significantly differently expressed in tumors from Black women in the VA-cohort compared to White women as a function of poverty burden. Linear regression coefficients were used to determine p-values using a T-test. Abbreviations: VA (Veterans Affairs cohort). 
