## Supplementary figures and images for "The impact of patient biology on racial disparities in breast cancer outcome"

### Figure S1

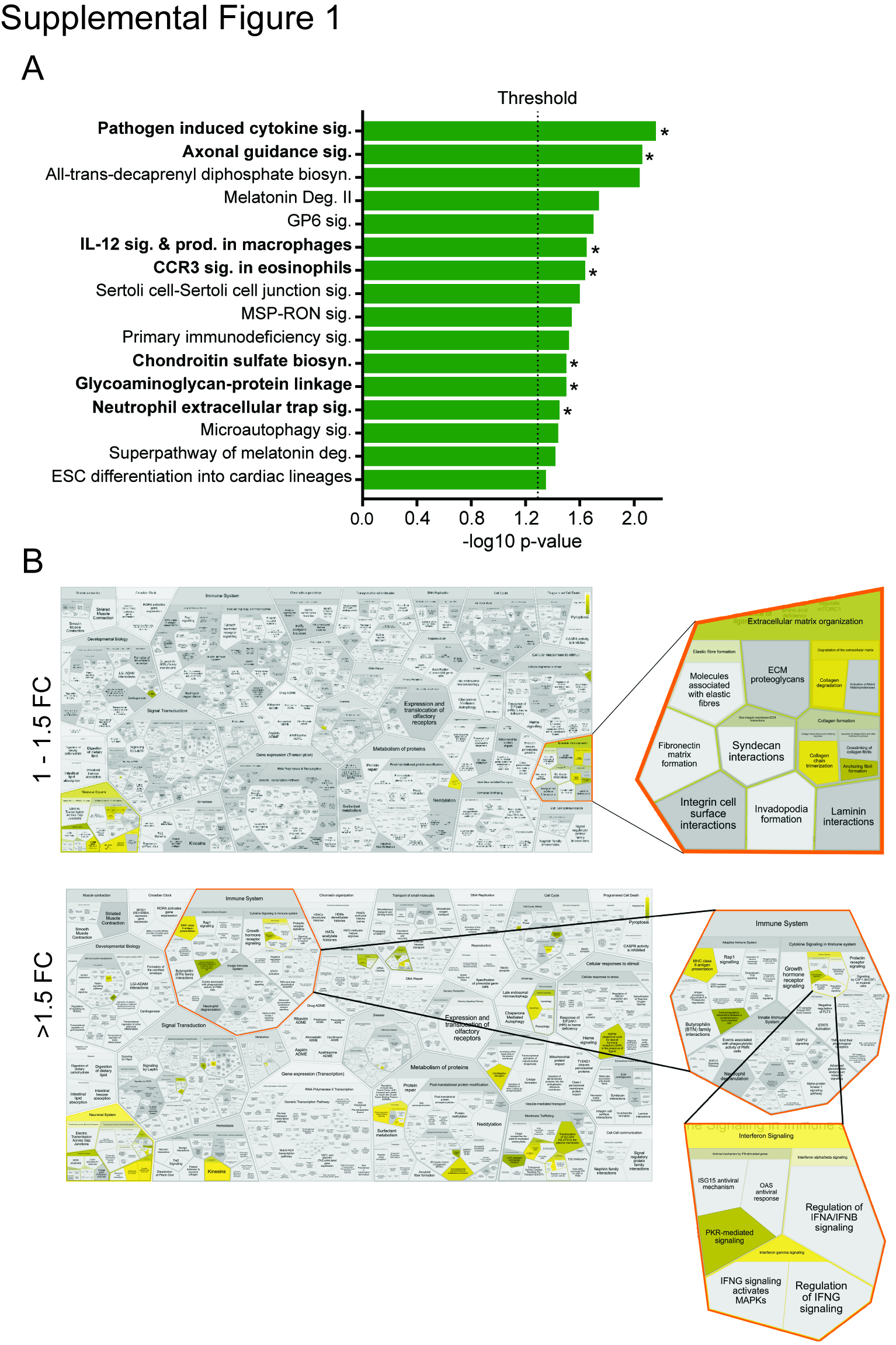

### Figure S2

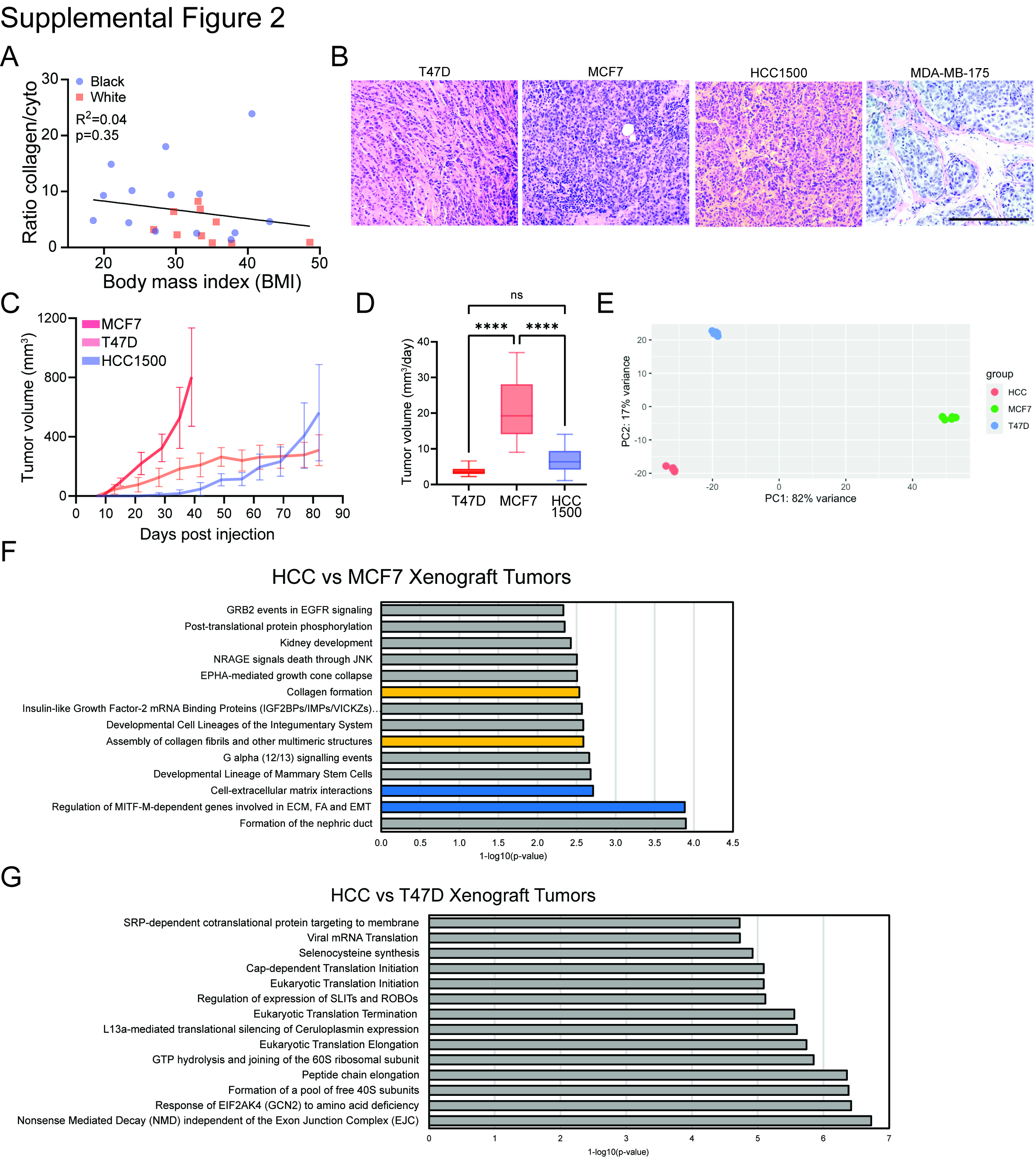

### Figure S3

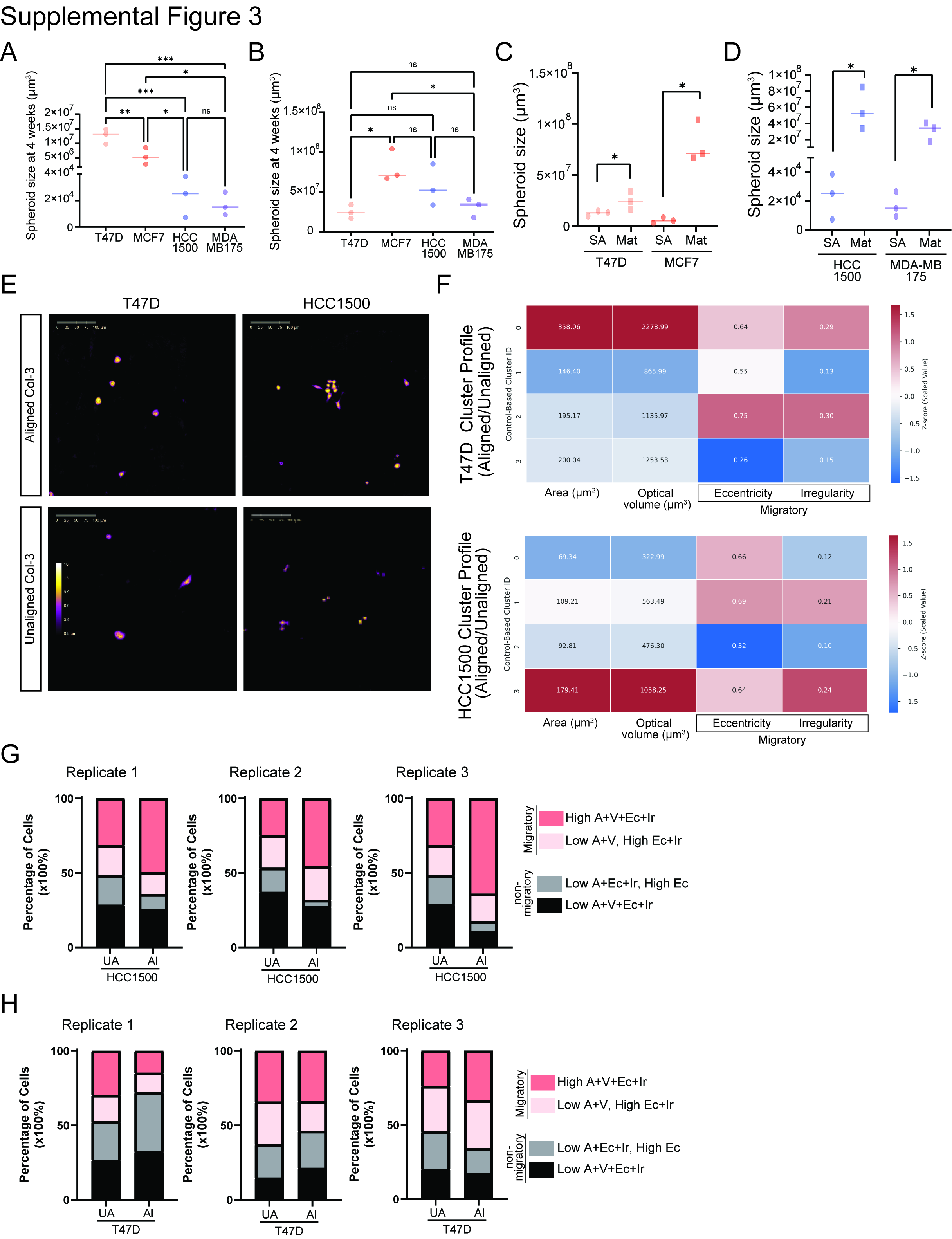

### Figure S4

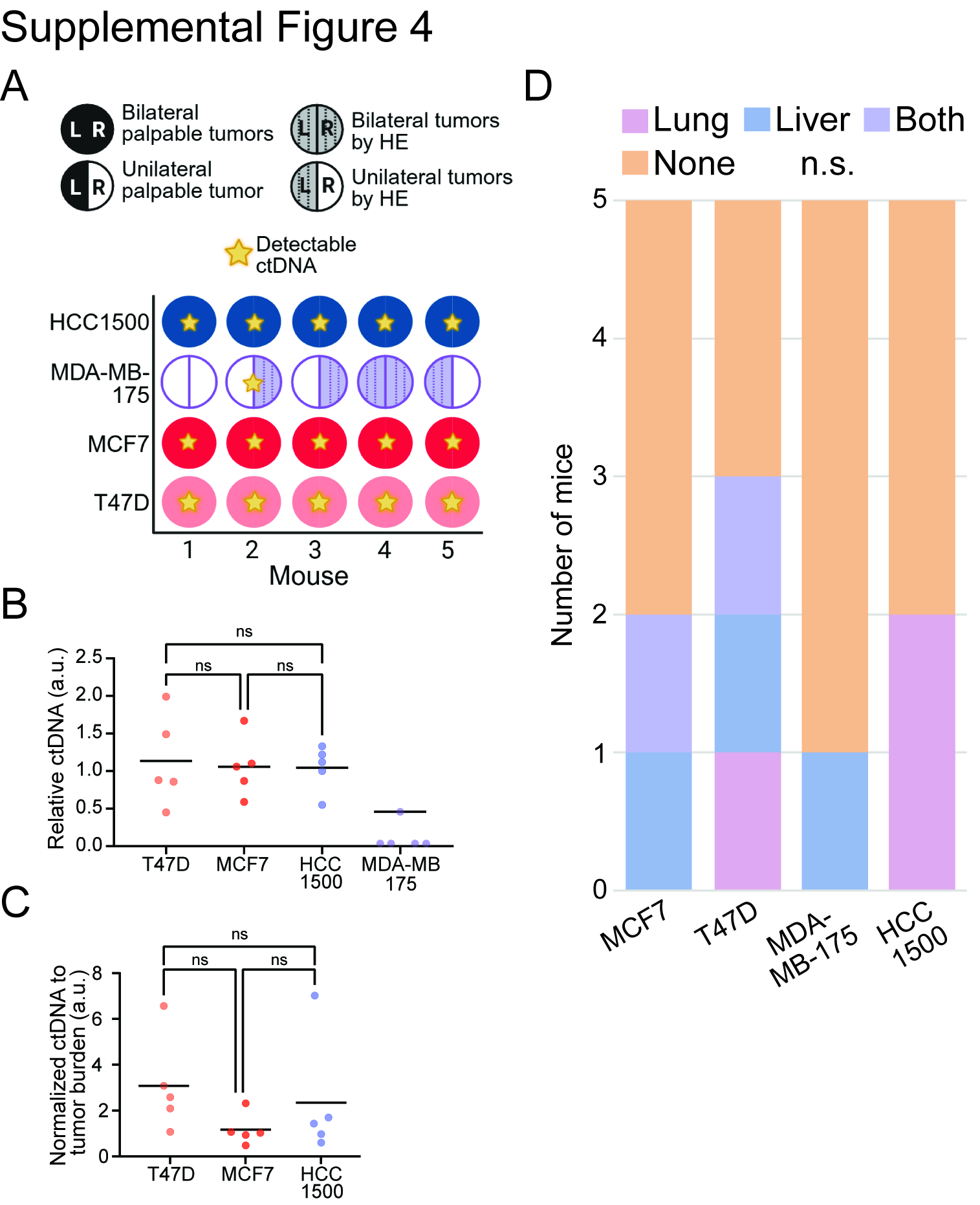

### Figure S5

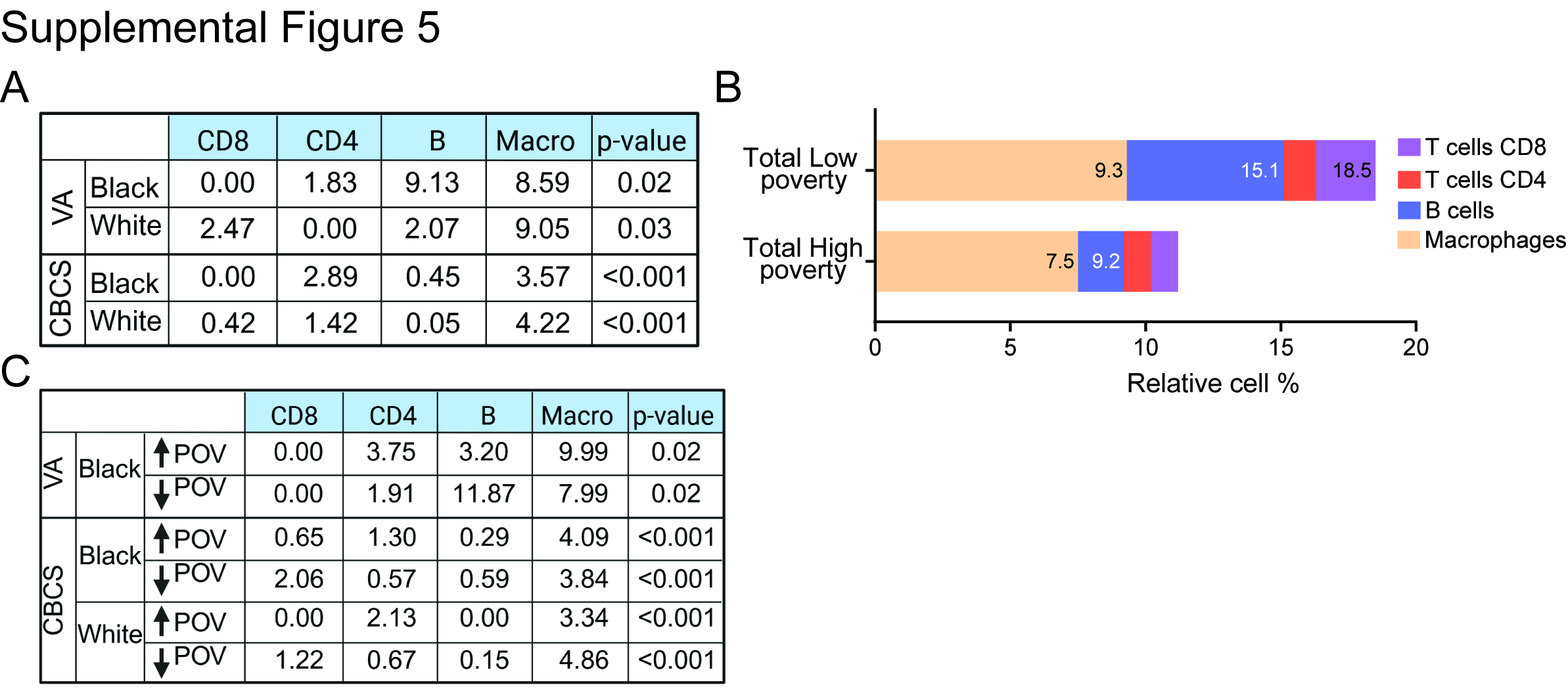

### Figure S6

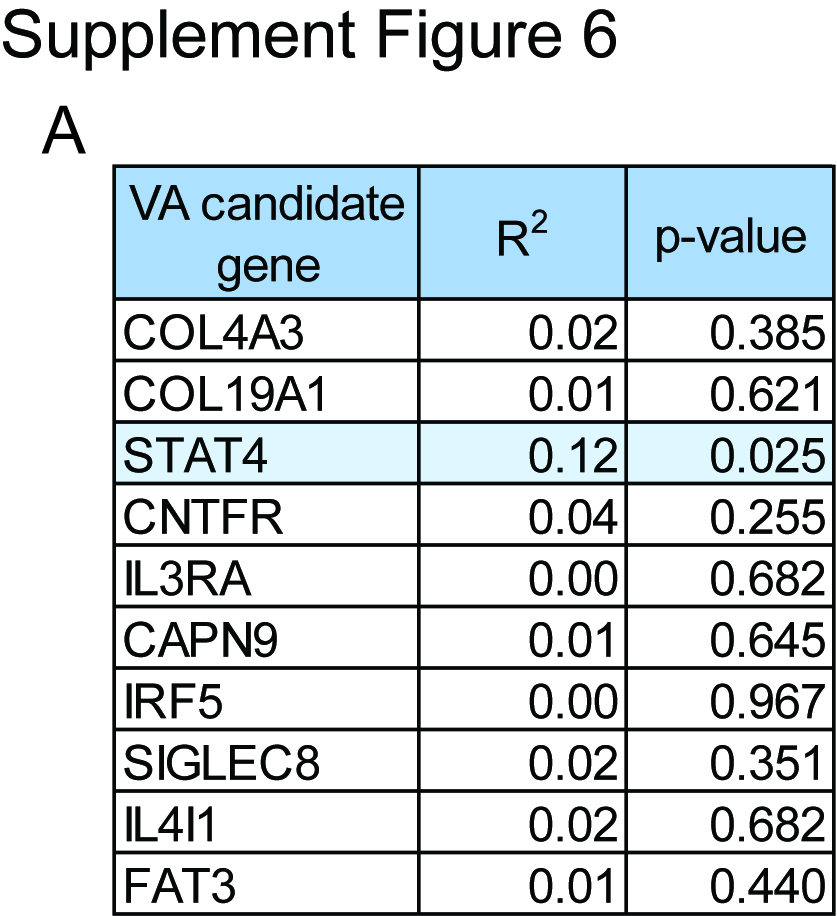
